# An Ovol1 Variant in Endothelial Cells Regulates Slug and is Essential for Pathological Angiogenesis

**DOI:** 10.64898/2026.10.31.763143

**Authors:** Kapil Thapa, Brandon Burow, Melody C. Baddoo, Marshall J. Peelle, Haiden Citron, K C Rajan, Shawn Sutherland, Christopher J. Hatch, Nadia Crawford, Ava Karp, Stryder M. Meadows, Daniel P. Howsmon, Brent A. Stanfield, Kelly Subramanian, Christopher C.W. Hughes, Jennifer S. Fang

**Affiliations:** Department of Cell and Molecular Biology, School of Science and Engineering, Tulane University, New Orleans, LA 70118; Tulane Cancer Center, School of Medicine, Tulane University, New Orleans, LA 70120; Department of Molecular Biology and Biochemistry, School of Biological Sciences, University of California-Irvine, Irvine, CA 92697; Department of Pathobiological Sciences, Louisiana State University-School of Veterinary Medicine, Baton Rouge, LA 70803; Department of Chemical and Biomolecular Engineering, School of Science and Engineering, Tulane University, New Orleans, LA 70118; Department of Biomedical Engineering, School of Biological Sciences, University of California-Irvine, Irvine, CA 92697; Department of Physiology, School of Medicine, Tulane University, New Orleans, LA 70120; Louisiana Cancer Research Center, New Orleans, LA 70124

**Author notes:** **Corresponding Author:** Jennifer S. Fang, 6400 Freret Street, 405 Paul Hall, New Orleans, LA 70118. Contributed Equally (Nadia Crawford, Ava Karp).

**Keywords:** Ovol1, Slug, Angiogenesis, Endothelial-to-Mesenchymal Transition, Cancer, Tumor Angiogenesis

## Abstract

During development, the blood vasculature expands by sprouting angiogenesis, but the programs that guide this process can become dysregulated in the pathological angiogenesis of neovascular diseases leading to excessive disorganized blood vessels. We previously reported that in endothelial cells, the pro-mesenchymal transcription factor Slug critically regulates angiogenesis through a partial and reversible endothelial-to-mesenchymal transition (pEndoMT); however, the upstream regulators that activate Slug and may contribute to pathological activation of the pEndoMT pathway in disease are still poorly understood. Here, we show that endothelial cells express an unusual low molecular weight variant of Ovol1 – a zinc-finger transcription factor that regulates mesenchymal transition in epithelial cells. Ovol1 and Slug are co-expressed in angiogenic endothelial cells, and using ChIP, we confirm direct Ovol1 binding to the endogenous Slug gene at a species-conserved binding motif. Whereas Ovol1 is typically a transcriptional repressor in epithelial cells, we find using a combination of silencing RNA and lentiviral overexpression constructs that the endothelial Ovol1 variant upregulates Slug, and is sufficient to drive pEndoMT marker expression and enhance endothelial cell migratory capacity. Silencing of Ovol1 abolished sprouting angiogenesis in a 3D *in vitro* sprouting angiogenesis assay, whereas constitutive overexpression of Ovol1 produced overaggressive and disorganized hypersprouting that depends upon Slug. We generated Ovol1 or Slug endothelial-specific knockout mice (Ovol1^ECKO^ and Slug^ECKO^) and found that healthy developmental angiogenesis of the neonatal mouse retina proceeds relatively normally in knockout animals. By contrast, Ovol1 was reduced in established post-natal blood vessels but became strongly re-expressed in tumor endothelial cells, and we found that colorectal cancer syngraft growth was significantly reduced in both Ovol1^ECKO^ and Slug^ECKO^ mice alongside a failure of tumor angiogenesis. Lastly, neovascularization in the oxygen-induced retinopathy (OIR) model was also reduced in Slug^ECKO^ mice. In conclusion, we report that an endothelial Ovol1 variant upregulates Slug and drives a mesenchymal-like state, and that this Ovol1-Slug pEndoMT signaling axis is essential for pathological angiogenesis in neovascular disease models.

**Statements and Declarations:** The authors declare that they have no financial or non-financial interests that are directly or indirectly related to this work.

## Introduction

Sprouting angiogenesis is a fundamental step of vascular development and post-natal blood vessel repair. Aberrant reactivation of developmental angiogenesis programs is believed to drive the overaggressive and disorganized pathological angiogenesis that occurs with neovascular diseases such as cancer and retinopathy.[1,2] However, the angiogenesis pathways that become aberrantly activated during pathological angiogenesis and that may contribute to disease progression are still incompletely understood.

During angiogenesis, endothelial cells (EC) – the cells that line the inner lumen of blood vessels – partially detach from the vessel wall and adopt highly characteristic migratory (i.e., “tip”) and proliferative (i.e., “stalk”) cell morphologies to form a new vessel sprout.[3] We previously showed that Slug – a member of the Snail/Gfi-1 (SNAG) family of zinc-finger transcription factors[4] – critically drives sprouting angiogenesis by enabling a **<u>p</u>**artial and reversible **<u>endo</u>**thelial-to-**<u>m</u>**esenchymal **<u>t</u>**ransition (pEndoMT) that temporarily decreases EC identity to enable transient acquisition of mesenchymal-like marker expression and cell behaviors (e.g., proliferation, migration, etc).[5,6] Importantly, abnormal activation of pEndoMT appears to be critical for diseased angiogenesis: Yang and colleagues recently reported that a significant proportion of tumor endothelial cells in a glioblastoma model exhibit a mesenchymal-like phenotype, and we and others find that individual pEndoMT master transcription factors – including Slug, Snail, and Twist – are essential for cancer angiogenesis.[7,2,8]. Yet, the upstream regulators that might contribute to aberrant activation of pEndoMT master transcription factors such as Slug are still largely unknown.

In epithelial cells, the Ovol1 zinc-finger transcription factor regulates epithelial-to-mesenchymal transition (EMT) by inhibiting pro-mesenchymal TGF-β signaling[9], suppressing transcription of pro-mesenchymal effectors, [10,11] [12] and promoting epithelial cell commitment.[13] Indeed, a recent ChIP-Seq analysis by Chen and colleagues revealed ∼1,100 direct Ovol1 target genes in epithelial cells,[14] underscoring a central role for Ovol1 in transcriptomic regulation of the mesenchymal transcription gene expression program. Interestingly, the phenotype of global Ovol1 knockout animals is strain-dependent: Ovol1 ablation is perinatally lethal in a C57/Bl6 background, but mixed 129Sv;C57Bl/6 Ovol1 knockout animals survive with pronounced defects in overall organ health[12] suggesting strain-specific compensation for Ovol1 loss during embryonic development. Failure of Ovol1 global-knockout animals to develop normally has largely been attributed to Ovol1’s function to limit EMT in epithelial cells. Meanwhile, a role for Ovol1 in endothelial cells has not yet been described. Therefore, in this study we set out to test the hypothesis that Ovol1 also plays a key role in endothelial cells to regulate Slug and pEndoMT in angiogenesis.

## Materials and Methods

### Cells

Primary human umbilical vein endothelial cells (HUVEC) (Sigma #200-05N) were cultured in EBM2 basal media (Lonza #CC-3156) supplemented with EGM2 BulletKit (Lonza CC-#4176). Primary normal human lung fibroblasts (Lonza # CC-2512) were cultured either in Dulbecco’s Modified Eagle Medium (DMEM) (Corning #10-013-CV) with 10% fetal bovine serum (FBS) (Corning #35010CV), or in FGM-2 (Lonza #CC-3132). 293T (ATCC #CRL-3216) and CMT93 cells (ATCC #CCL-223) were also cultured in DMEM supplemented with 10% FBS. MC38 cells (Kerafast #ENH204-FP) were cultured in DMEM supplemented with 10% FBS and 0.1mM non-essential amino acids. SW620 (ATCC #CCL-227) and YCCEL1 (gift of Dr. Erik Flemington, Tulane) were cultured in RPMI 1640 medium (Cytvia #SH30027.02) supplemented with 10% FBS. All cells were cultured in 5% CO_2_ at 37°C and primary cells were used for Bead Assay experiments no later than passage 6, or no later than passage 8 for monoculture studies.

### Lentivirus

293T cells were transfected using Lipofectamine 2000 (Invitrogen #52887) with 3^rd^ generation lentiviral packaging plasmids mDLG/pRRE (Addgene #12251), MD2.G (Addgene #12259), RSV-Rev (Addgene 12253), and lentiviral expression constructs (i.e., Ovol1 ORF: Addgene #144737[15], eGFP: Addgene #36083). Culture supernatant was collected on days 2-4, and lentivirus was concentrated using 50% polyethylene glycol 8000 (Promega #V3011) for 48h at 4°C prior to collection by centrifugation and resuspension in PBS. Cells were transduced with lentivirus in the presence of 10μg/ml polybrene (Santa Cruz #NC9840454) at MOI ≈ 1.

### Silencing RNA

Cells were plated at sub-confluent density overnight, and culture media was refreshed 30 minutes prior to transfection. 25nM of SMARTpool silencing RNA (si-Ovol1: Dharmacon #L-006543-01-0005; si-Slug: Dharmacon #L-017386-00-00051) or non-targeting control (Dharmacon #D-001810-10-2) was transfected according to manufacturer’s protocols using Lipofectamine RNAiMAX (Invitrogen #13778150) and cultured overnight in standard growth media. Culture media was replaced at 16h post-transfection, and cells were harvested at 48 hours for subsequent experimentation or analysis.

### Fibrin Gel Bead Assay

To model sprouting angiogenesis *in vitro*, the Fibrin Gel Bead Assay was performed as previously described.[16] In brief, SphereCol beads (Advanced Biomatrix #5138) were coated with HUVEC over-night and embedded in 2.5mg/mL fibrin (MilliporeSigma #341573) hydrogel containing 60,000 cells/mL lung fibroblasts. The resulting cell suspension was mixed with 0.625U bovine thrombin (Sigma #T4648) at a concentration of 500 beads/mL. Approximately 250 beads were embedded per well of a 24-well plate, and an additional 30,000 fibroblasts were seeded on top of the fibrin gel. Beads were fed with complete EGM2 media every other day and cultured for up to 10 days.

### Immunocytochemistry

For adherent cells, cells cultured on gelatinized glass coverslips were fixed with 4% paraformaldehyde (PFA) (Sigma #47608) prior to blocking with 3% bovine serum albumin (BSA)(Fisher Scientific #BP9703-100) 0.1% Triton X 100 (Thermo #A16046) in 1X PBS, and subsequent primary antibody incubation over-night at 4°C rocking in 1% BSA 0.1% Triton X 100 in PBS. Following primary antibody incubation, coverslips were washed and incubated in fluorescently-conjugated secondary antibody for 2 hours at room temperature. Coverslips were counterstained with Hoechst (1μg/mL, Sigma #B2261) for 15 minutes, RT. Coverslips were mounted onto glass slides with Vectashield mounting media (Fisher #NC9265087) and imaged using Leica DMi8 or Nikon ECLIPSE Ti2. For Bead Assay, fibroblasts cultured on hydrogel surface were removed by trypsinization prior to bead fixation by addition of 2% PFA for 10 minutes RT. Cultures were permeabilized with 0.5% TX100 1X PBS for 45 minutes RT and blocked with 1% BSA 5% FBS 0.1% Tween-20 (Sigma #P9416), following by incubation in primary antibodies for 24 hours at 4°C. Following wash, cultures were incubated in fluorescently-conjugated secondary antibodies for 2 hours at room temperature. After wash, cultures were counterstained with Hoechst in 1X PBS for 15 minutes, RT and imaged with Nikon ECLIPSE Ti2.

### Antibodies

For immunofluorescence: Goat anti-CD31 (1:200, R&D #AF3628); Rabbit anti-Ovol1 (1:500, Proteintech #14082-1-AP); Mouse anti-Slug (1:300, Santa Cruz #sc-166476); Mouse anti-αSMA (1:500, Sigma #A5228); Rabbit anti-ERG Alexa fluor 647 (1:200, Abcam #ab196149); Donkey anti-Goat Alexa fluor 647 (1:200 - 1:600, Invitrogen #A21447); Donkey anti-rabbit Alexa fluor 594 (1:200 - 1:600, Invitrogen #A21207); Donkey anti-mouse Alexa fluor 488 (1:200 - 1:600, Invitrogen #A21202). For immunohistochemistry: Rabbit anti-Ovol1 (1:100, Atlas #HPA003984). For Western blot: Rabbit anti-Ovol1 (1:500, Proteintech #14082-1-AP), Mouse anti-V5 (1:250, Biorad # MCA1360GA), Rabbit anti-αTubulin (1:1000, Invitrogen #PA5-29444); Goat anti-GAPDH (1:1000, Novus #NB300-320); Goat IR Dye 800 CW Goat anti-rabbit (1:5000, LICORbio #926-32211); Donkey IR Dye 800 CW anti-goat (1:5000, LICORbio #926-32214).

### Chromatin-immunoprecipitation (ChIP)

HUVEC monocultures were resuspended in PBS at 5 x 10^6^ cells/mL and crosslinked with 1% PFA, followed by glycine wash. Samples were treated with protease inhibitors (1mM PMSF, MilliporeSigma #P7626-1G and 1X protease inhibitor cocktail, Roche #11697498001), and chromatin was prepared using a Zymo-Spin ChIP kit (Zymo #D5209-A) by manufacturer’s protocol. Chromatin was sheared using a Q55 Sonicator (QSonica #Q55-110) on ice for 4 cycles (30 sec “ON”, 30 sec “OFF” at 40% amplitude). We subsequently confirmed fragmentation of total chromatin to ∼100bp-500bp fragments via gel electrophoresis. Sheared chromatin was incubated with Ovol1 antibody (4μg, Proteintech #14082-1-AP) or isotype control (Rabbit IgG, 4μg, Invitrogen #31235) over-night at 4°C, rocking. Reverse crosslinking was performed in 5M NaCl at 75°C for 5 minutes, and at 65°C for 30 minutes, prior to ChIP DNA purification using Zymo-Spin ChIP kit. Purified ChiP DNA was analyzed by quantitative PCR for pull-down of the Slug gene using primers listed in **Table 1**. A 2% fraction of starting chromatin sample was used as input control, and ChiP DNA enrichment was graphed as % of input

**Table 1.**
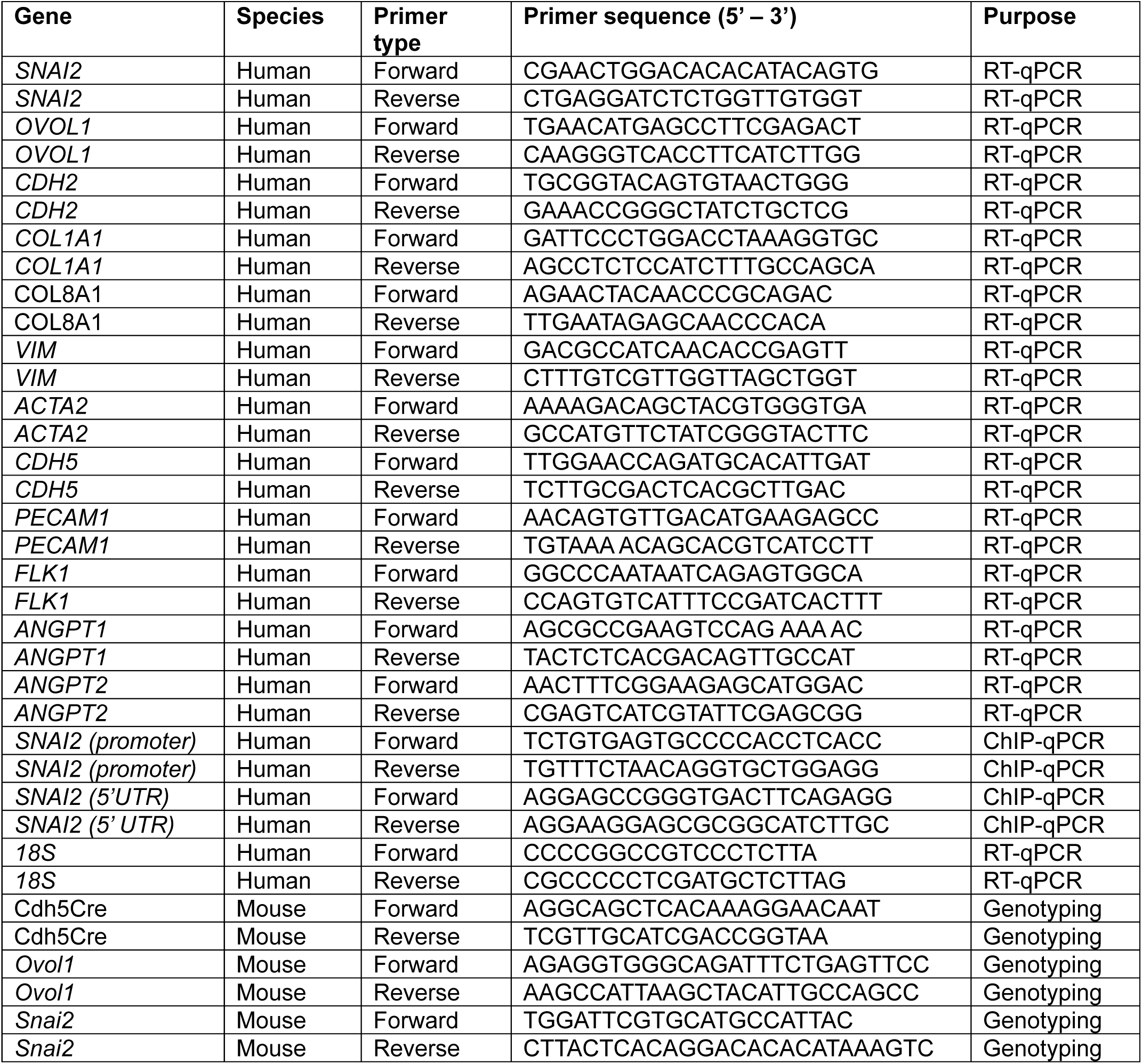
Primer Sequences.

### Quantitative PCR

RNA was extracted using a Quick-RNA Microprep kit (Zymo #R1051), and quality and concentration of sample was confirmed using a Nanodrop 2000 (Thermo). cDNA synthesis was performed using the iScript gDNA Clear cDNA Synthesis Kit (Biorad 1725035BUN) according to manufacturer’s protocol. Resulting cDNA libraries were probed by quantitative PCR using PerfeCTa SYBR Green Supermix (Quantabio #95054-500) and gene-specific primers (**Table 1**) using a CFX96 Touch Real-Time PCR Detection System (Biorad).

### Western Blot

Cells were lysed in RIPA buffer (Thermo #89900) supplemented with protease inhibitors cocktail (Roche #11697498001), PMSF (Sigma #P7626), and sodium orthovanadate (Sigma #S6508). Lysates were maximally sonicated three times for 30 seconds each cycle. Protein concentration was measured using a BCA protein assay kit (Thermo #A55860). 30μg - 50μg of protein was loaded and run in a precast gel (Biorad #4561084) and transferred to PVDF membranes (Biorad #10026934) for 2 hrs. Membranes were blocked with 5% non-fat dry milk (NFDM) milk in Tris Buffered Saline (TBS) for 1 hour at room temperature. Primary antibodies were diluted in 1% NFDM milk in TBS and incubated at 4C overnight, while secondary antibodies were diluted similarly and incubated at room temperature for 2 hours. Membranes were imaged with LiCor Odyssey Imaging System 9120.

### Next-Generation Sequencing and Analysis

RNASeq was performed in collaboration with the Louisiana Cancer Research Center’s Translational Genomics Core. In this study, three replicates were generated per experimental or control group. RNA was harvested using Quick-RNA Micro-prep kit (Zymo #R1051), and concentration was determined using a Qubit fluorometer. cDNA libraries were generated using NextSeq 1000/2000 P2 Reagents (200 Cycles) v3 kit (Illumina #20046812). For quality control, the DNA 1000 kit (Agilent #5067-1505) was used with Agilent 2100 bioanalyzer to determine cDNA fragment size. cDNA concentration was measured with Qubit dsDNA quantification assay kit (Invitrogen #Q32851). Samples were pooled and sequenced using an Illumina NextSeq 1000/2000. Pre-processing and alignment were performed using Tulane’s Cypress high-performance computing cluster. In brief, quality control was done by fastp (v.0.23.4)[17] with default parameters, and aligned using the Ensembl GRCh38 human reference genome using STAR (v.2.5.2a)[18] and processed further with samtools (v.1.10). Feature-count matrix was acquired by running featureCounts() function from Rsubread (v 2.16.1). All downstream analysis was done in R (v4.3.2).[19] R-package org.Hs.eg.db (v3.17.0) was used to convert entrez IDs to gene symbols. Differential analysis was done using DESeq2 (v1.40.2). Pathway analysis was performed using clusterProfiler (v4.8.3) or PANTHER (v.19.0).[20] Top pathways that were significantly enriched (p_adjust_ < 0.05) were selected and visualized with dotplot. Gene interaction networks were generated using the STRING database platform, with cluster analysis performed based on node density.

Single-nucleus RNA and ATAC-seq data of tumor and adjacent healthy tissue from nine colorectal cancer patients were obtained from the published data deposited by Yu and colleagues[21] (GEO accession GSE294559) as 10x Genomics Multiome Cell Ranger ARC outputs. The data included feature-barcode H5 files and corresponding ATAC fragment files. Only normal (healthy) colorectal tissue was considered for this analysis. Multiome objects containing RNA and ATAC assays were generated and processed using Seurat (v5.2.1) and Signac (1.14.0) with the hg38 reference genome and gene annotations derived from EnsDb.Hsapiens.v86. Per-nucleus quality control included RNA feature counts, transcription start site enrichment nucleosome signal and reads-based fraction of reads in peaks (FRiP). Nuclei were retained with 500-6000 detected RNA features, TSS enrichment >2, nucleosome signal <2, and FRiP >3. Following QC, RNA counts were log-normalized, highly variable genes were identified and scaled, and principal component analysis was performed. ATAC data was normalized using Term Frequency-Inverse Documents Frequency (TF-IDF) followed by latent semantic indexing (LSI). RNA and ATAC were integrated using Seurat’ s weighted nearest-neighbor framework, followed by joint UMAP dimensionality reduction and clustering. Cell populations were annotated at the WNN-cluster level using SingleR with the Human Primary Cell Atlas (HPCA) and Blueprint/ENCODE reference datasets, followed by manual review using cluster-specific marker genes. Differential RNA expression and chromatin accessibility were assessed using patient-level pseudobulk counts with DESeq2 (v1.42.1), incorporating patient as a blocking factor.

### Mice

For Ovol1^ECKO^ mice, Ovol1^tm1a(KOMP/Wtsi)^ mice (i.e., Ovol1^flox/flox^) – in which a cassette containing *Frt*, *LacZ*, and loxP sites are inserted into the endogenous Ovol1 locus[22,23] – were crossed with mice expressing an endothelial-specific (via *Cdh5* (VE-Cadherin) promoter) Cre recombinase[24] to generate endothelial-specific Ovol1 knockout mice (i.e., Ovol1^ECKO^). For Slug^ECKO^ mice, we partnered with the University of California-Irvine Transgenic Mouse Facility to use CRISPR/Cas9 gene editing to insert flanking loxP sites around the endogenous *Snai2* gene locus of C57/Bl6 mice. The presence of flanking loxP inserts were confirmed by PCR genotyping (primers listed in **Table 1**), and mice were bred to Slug^flox/flox^ homozygosity before being crossed with the Cdh5Cre driver mouse line.[24] Slug^ECKO^ and Ovol1^ECKO^ mouse lines were subsequently transferred to Tulane University for further study. Ovol1^ECKO^ and Slug^ECKO^ mice were maintained heterozygous for the Cre transgene and homozygous for flanking loxP sites, and all mice were biopsied and PCR genotyped at time of weaning using primers listed in **Table 1**. In all experiments, Cre-positive homozygous floxed (i.e., knockout) mice were compared against their Cre-negative homozygous floxed littermate controls, and equal ratios of male and female mice were included in each control or experimental group. For xenograft experiments, NOD.Cg-Prkdcscid Il2rgtm1Wjl/SzJ (NSG) mice (Jackson Laboratories # 005557) were housed at the Louisiana State University School of Veterinary Medicine Division of Laboratory Animal Medicine. All animal work was performed in accordance with the University of California-Irvine, Tulane University, and Louisiana State University A&M College Institutional Animal Care and Use Committee policies and oversight.

### FACS

Post-natal day 15 mice were euthanized and lung tissue was digested in 3mg/mL collagenase type I (Thermo #17100017) for 1hr at 37°C to generate a single-cell suspension. Cells were incubated in the presence of fluorescently-conjugated antibodies for CD31 (1:200, BD Biosciences #551262) and CD45 (1:200, BD Biosciences #553088). Labelled cells were sent to the Flow Cytometry and Cell Sorting Facility, Department of Microbiology, Tulane University where ∼500,000 CD31+/CD45-endothelial cells were sorted per mouse. Genomic DNA was isolated from sorted lung EC using a Zymo Quick-DNA Miniprep kit (Zymo #D3024) according to manufacturer’s protocol.

### Neonatal Retina and Oxygen-Induced Retinopathy Model

Neonatal retina vasculature was harvested and analyzed as previously described.[25] In brief, mice were euthanized at post-natal day 6, followed by enucleation, tissue fixation in 4% PFA, and dissection. Following blocking in 5% BSA 0.3% Triton X 100 (i.e. TX100) in PBS for overnight, retinas were incubated in primary antibody diluted in 1% BSA 0.3% TX100 PBS for 24 hours at 4°C rocking. For Ovol1 staining, retinas were post-fixed with 4% paraformaldehyde (PFA) for 3 hours at room temperature, dehydrated (50% methanol, 5 minutes) and methanol fixed overnight. Following methanol rehydration, retinas underwent antigen retrieval (10µg/ml Proteinase K (Thermo #EO0491), 2 minutes) followed by incubation in blocking solution (10% donkey serum Sigma #D9663, 1% DMSO, 0.5% TX100 PBS) for 1 hour at room temperature. Retinas were subsequently incubated in staining solution containing primary antibodies for 24 hours at 4°C, rocking. Following wash, retinas were incubated in fluorescently-conjugated secondary antibody and fluorescently-conjugated isolectin B4 (Invitrogen #I21411) diluted in 1% BSA 0.25% TX100 in PBLEC buffer[25] or PBS for 2 hours at room temperature or overnight at 4°C, rocking. Following wash, nuclei were counterstained with Hoechst 33342 (1μg/mL, Sigma #B2261) 15 minutes, room temperature or with anti-ERG Alexa Fluor 647 for 2 hours room temperature. Stained retinas were mounted on glass slides in Vectashield and imaged by confocal microscopy using a Nikon ECLIPSE Ti2. For oxygen-induced retinopathy (OIR) experiments, pups and nursing dams were placed at P7 in hyperoxic (75% O_2_) conditions for five days, followed by return to room air (20% O_2_) at P12 for five days to simulate hypoxia, as previously described.[26] After this exposure, pups were euthanized at P17 and tissue was harvested and processed as described above.

### Tumor Syngraft and Xenograft

CMT93 mouse colorectal cancer cells were suspended in Matrigel (Corning #CB-40234) at 2 x 10^6^ cells/mL. MC38 cells, another murine colon cancer cell line, were suspended in Matrigel at 0.5 - 2 x 10^6^ cells/mL. Appropriate volume of cell suspension (500 μl for CMT93; 200-500 μL for MC38) was subcutaneously injected into the dorsal right flank of 8 to 12-week-old mice. Tumor growth was monitored regularly for up to 6 weeks using digital calipers. For tumor harvest, animals were deeply anaesthetized by isoflurane and perfusion-fixed with 4% PFA. Tumors were surgically removed and post-fixed with 4% PFA over-night at 4°C, followed by serial dehydration in ethanol. Tumor tissue was briefly incubated in Histoclear (VWR #101412-878) to remove ethanol and embedded into paraffin for immunohistochemistry. For guinea pig tumor xenografts, 1 x 10^6^ 104C1 guinea pig embryonic fibroblasts (ATCC #CRL-1405) suspended in 100 μl of sterile saline was injected subcutaneously into 6- to 8-week-old female NSG mice. Tumor growth was monitored over a period of 50 days using digital calipers. Animals were euthanized when tumor volumes reached approximately 200 mm^3^ in volume. Tumors were excised at necropsy, fixed overnight in 10% neutral buffered formalin, and submitted to the Louisiana Animal Disease Diagnostic Laboratory (LADDL) histology core for paraffin embedding, sectioning, and immunohistochemistry as described below.

### Immunohistochemistry

Formalin-fixed paraffin-embedded tissue blocks were cut into 5 µm thick sections using a Thermo Fisher Scientific HM325 rotary microtome and adhered to glass slides. Slides were deparaffinized with xylene and rehydrated by serial ethanol incubation, followed by antigen retrieval in sodium citrate buffer (Sigma #71402) for 5 minutes at high temperature and pressure. Tissue sections were blocked in 5% donkey serum (Sigma #D9663) 0.1% TX100 PBS for 1 hour at room temperature, followed by incubation in primary antibody overnight at 4°C. Following wash, tissue sections were incubated in fluorescently-conjugated secondary antibody diluted in 0.1% TX100 PBS for 2 hours at room temperature. Following wash, nuclei were counterstained in Hoechst 33342 (1µg/mL, Sigma #B2261). Slides were mounted with glass coverslips in Vectashield and imaged with Nikon ECLIPSE Ti2.

### Vascular Morphometry

All images were analyzed in Fiji/ImageJ. For neonatal retina, vascular morphometry was performed as previously described.[25] In brief, vascular outgrowth was calculated as the relative distance from optic nerve head to vascular front vs. total distance from optic nerve head to retinal edge. Arterial outgrowth was calculated as relative length of αSMA^+^ arteries from optic nerve head to loss of signal vs. vascular outgrowth distance. For other morphometric analyses (e.g., vessel area), an 850µm X 700µm ROI was selected at the angiogenic front, encompassing the majority of the angiogenic front for each petal including distal regions of both veins and arteries. Angiotool 2.0 was used measure IB4^+^ vessel area for each ROI, and the mean vessel area was calculated per retina and averaged for each mouse. For tip cell number, a 1000µm X 500µm ROI was selected at the vascular front, and tip cells (i.e. EC with filipodia-like projections) were manually counted in ImageJ. The mean tip cell number for each mouse was calculated across multiple ROIs for each sampled retinas. For tumor syngrafts, 10-12 500µm X 500µm ROIs were randomly selected for each tissue section and microvascular density -- calculated as the number of CD31^+^ vessels per area of tumor -- was quantified for each ROI to obtain a mean density value for the tumor.

### Hematological Analysis

Immediately following euthanasia, blood was collected from 8-12 week old mice into EDTA-treated tubes (Greiner Bio-One #450474). Hematological profile was collected using a VETSCAN HM5 automated analyzer.

### Software

Alignments were performed using Clustal and other tools available through the UCSC Genome Browser.[27] Predicted protein molecular weights were calculated using ExPasy (https://web.expasy.org/compute_pi/) based on peptide sequences from Genbank. Protein structures were predicted using AlphaFold (http://alphafoldserver.com), compared using ChimeraX (UCSF) (https://www.cgl.ucsf.edu/chimerax/), and internal translational start sites were identified using NetStart 2.0[28] or ATGpr (https://atgpr.dbcls.jp/).

### Statistics

As indicated in figure legends, two-group comparisons were performed using Unpaired Students’ t-test. Multi-group comparisons were performed using 1-way or 2-way ANOVA with mixed effects and post hoc t-test adjusted for multiple comparisons. Non-parametric data was compared using Wilcoxon rank-sum test, or Chi square test to identify statistical differences in observed vs. expected frequency data. In all tests, α was set at 0.05.

## Results

### Ovol1 is expressed in endothelial cells during angiogenesis

Ovol1 has previously been shown to regulate the mesenchymal transition program in epithelial cell types,[13] but its expression and function in endothelial cells is largely unknown. To first determine if Ovol1 is endogenously expressed in EC, we performed Western blot analysis of primary human umbilical vein endothelial cells (HUVEC, 3 independent donors) vs. other cancer and non-cancer epithelial and fibroblast cell types, including primary normal human lung fibroblasts (2 independent donors), human embryonic kidney cells (293T, an immortalized cell line with epithelial-like morphology), a human gastric epithelial cancer cell line (YCCEL1), and a human colorectal adenocarcinoma cell line (SW620). We identified robust Ovol1 signal in all cell lines (**Figure 1a**, full-sized blots in **Supplementary Figure 1**). As expected, both epithelial cancer cell lines (i.e., YCCEL1, SW620) predominantly expressed an Ovol1 variant that migrates at ∼30kDa – consistent with the predicted molecular weight of 30.26kDa for full-length Ovol1 (Genbank NP_004552.2, predicted molecular weight via ExPasy). By contrast, endothelial cells and 293T predominantly express an Ovol1 variant that migrates at ∼25kDa, with no band observed at ∼30kDa. Fibroblasts endogenously express both high and low molecular weight Ovol1 variants. To independently confirm that endothelial cells endogenously express Ovol1, we also reanalyzed a published single-cell multiomic dataset that captures both endothelial and epithelial cell populations from healthy adult colon donor tissue.[21] In this dataset, we also identified detectible endogenous transcript for Ovol1 in endothelial cells of healthy post-natal tissue, although endothelial Ovol1 expression was substantially lower than in healthy colon epithelial cells (**Supplementary Figure 2**).

**Figure 1:**
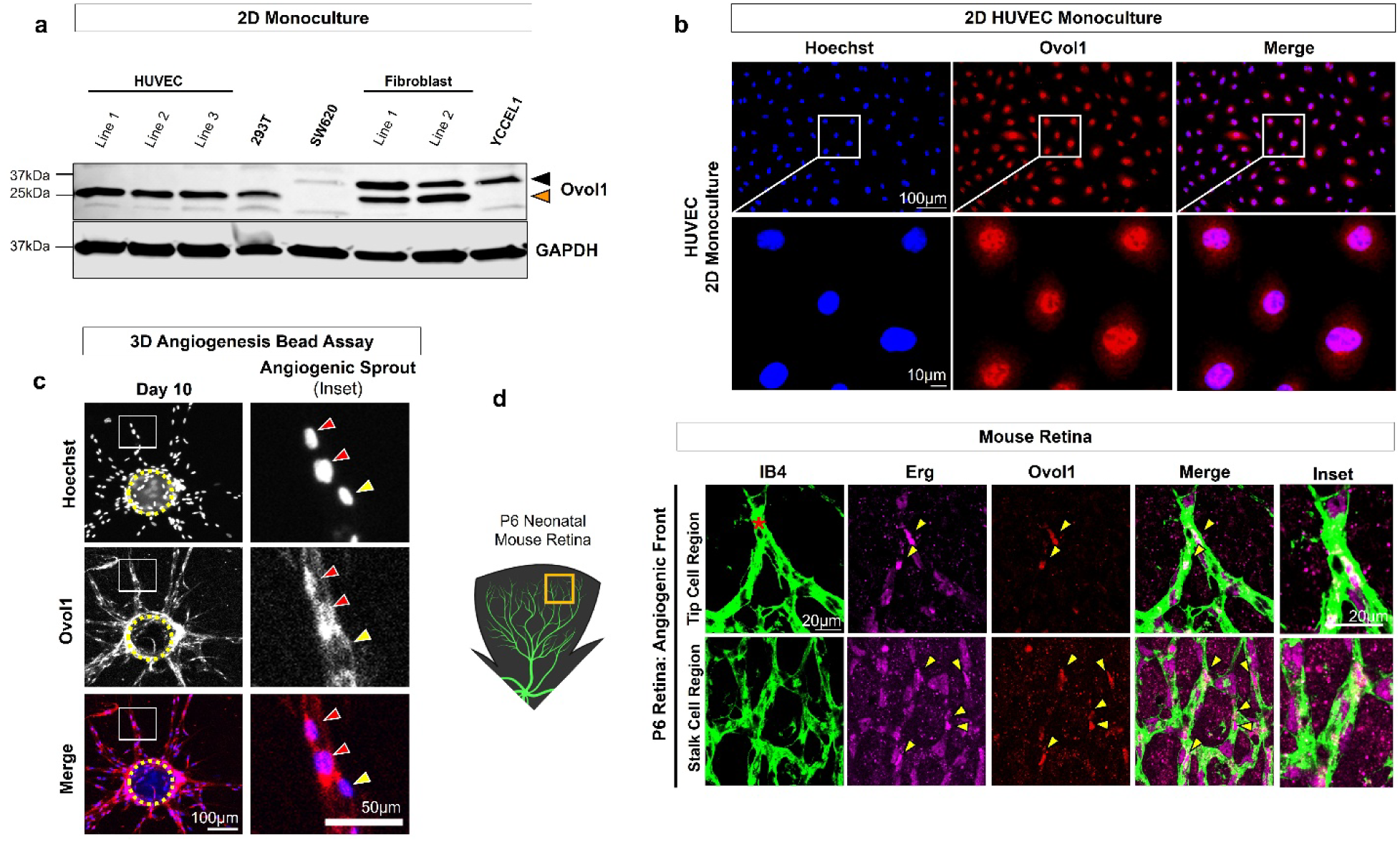
Ovol1 is expressed in endothelial cells during angiogenesis. **a)** Ovol1 is detected by Western blot in total protein lysate harvested from primary human umbilical vein endothelial cells (HUVEC, 3 independent donor lines), primary human normal lung fibroblasts (2 independent donor lines), immortalized epithelial-like human embryonic kidney (HEK293T) cells, a gastric epithelial cancer cell line (YCCEL1), and colorectal adenocarcinoma cancer cell line (SW680). All cells were grown in 2D monoculture. Cancer cells primarily express a ∼30kDa species of Ovol1 (black arrowhead), whereas in HUVEC, a ∼25kDa Ovol1 species (orange arrowhead) predominates. **b)** In HUVEC, Ovol1 immunofluorescent signal (red) is predominantly nuclear (Hoechst, blue). **c)** Ovol1 (red) is strongly detected in many (red arrowheads), but not all (yellow arrowhead), nuclei (Hoechst, blue) of angiogenic EC that form sprouts in the *in vitro* Bead Assay sprouting angiogenesis model at Day 10 (bead, dashed yellow circle). **d)** In the post-natal day 6 (P6) retina of wild-type mice, Ovol1 is expressed in endothelial cell nuclei (marked by endothelial-specific nuclear marker, Erg) at the angiogenic front of the developing retinal vasculature (marked by endothelial marker isolectin B4 (IB4)), including in EC at the tip cell position (red asterisk, *).

Ovol1 is a zinc finger transcription factor that becomes activated to translocate into the nucleus and regulate downstream gene expression.[29] Using immunofluorescence, we found Ovol1 to be predominantly nuclear in HUVEC cultured in (growth factor-rich) complete EC media under 2D monoculture conditions (**Figure 1b**). To further assess endothelial Ovol1 expression and subcellular localization during sprouting angiogenesis, we leveraged the 3D Fibrin Gel Bead Assay (i.e., “Bead Assay”) *in vitro* model of sprouting angiogenesis,[16] wherein HUVEC are cultured on the surface of collagen-coated beads embedded in a 3D fibrin gel in the presence of fibroblasts. Under these conditions, HUVEC form 3D angiogenic sprouts over the course of 10 days. We harvested beads at Day 10 and performed immunofluorescence, where we observed that Ovol1 was expressed in the nucleus and perinuclear space of some EC that comprise an angiogenic sprout (**Figure 1c**, red arrowheads) but is not significantly expressed in others (**Figure 1c**, yellow arrowheads). To further confirm the physiological relevance of these results, we turned to the neonatal mouse retina in which a superficial vasculature forms on the inner retinal surface between post-natal (P) days 0-9 via sprouting angiogenesis at the expanding network front with subsequent remodeling to support vessel maturation closer to the optic nerve head. In wild-type P6 retinas, we observed numerous Ovol1^+^ EC nuclei at the angiogenic front, including in EC occupying both tip and stalk cell positions (**Figure 1d**).

### Ovol1 regulates angiogenic signaling pathways

To identify signaling pathways regulated by Ovol1 in endothelial cells, we performed bulk RNA sequencing on EC transfected with silencing RNA to eliminate endogenous Ovol1 expression. In HUVEC, si-Ovol1 achieved ∼50% knockdown of endogenous Ovol1 mRNA (**Supplementary Figure 3a**) and almost completely abolished the ∼25kDa Ovol1 band by Western blot (**Supplementary Figure 3b**).

Transcriptomic analysis of si-Ovol1 EC (vs. si-Ctrl) revealed 1,108 differentially-expressed genes (adjusted p-value < 0.05), including 467 upregulated and 641 downregulated genes following Ovol1 silencing (**Figure 2a**, DEG list in **Supplementary Table 1**). Interaction network analysis of these Ovol1-dependent DEGs revealed 20 cell signaling clusters, with the largest two associated with cell proliferation and cell growth signaling pathways (**Supplementary Figure 4**). Gene Ontology (GO) analysis of these DEGs revealed 518 distinct GO terms (**Supplementary Table 2**), including terms related to angiogenesis, vascular development, and blood vessel function (**Figure 2b**). These angiogenesis-associated GO terms were driven by differential expression of 102 angiogenesis-related genes (**Figure 2c, Supplementary Table 3**), representing ∼10% of all Ovol1-dependent DEGs in our dataset. Taken together, these data support Ovol1 as a critical regulator of angiogenesis and vessel growth in endothelial cells.

**Figure 2.**
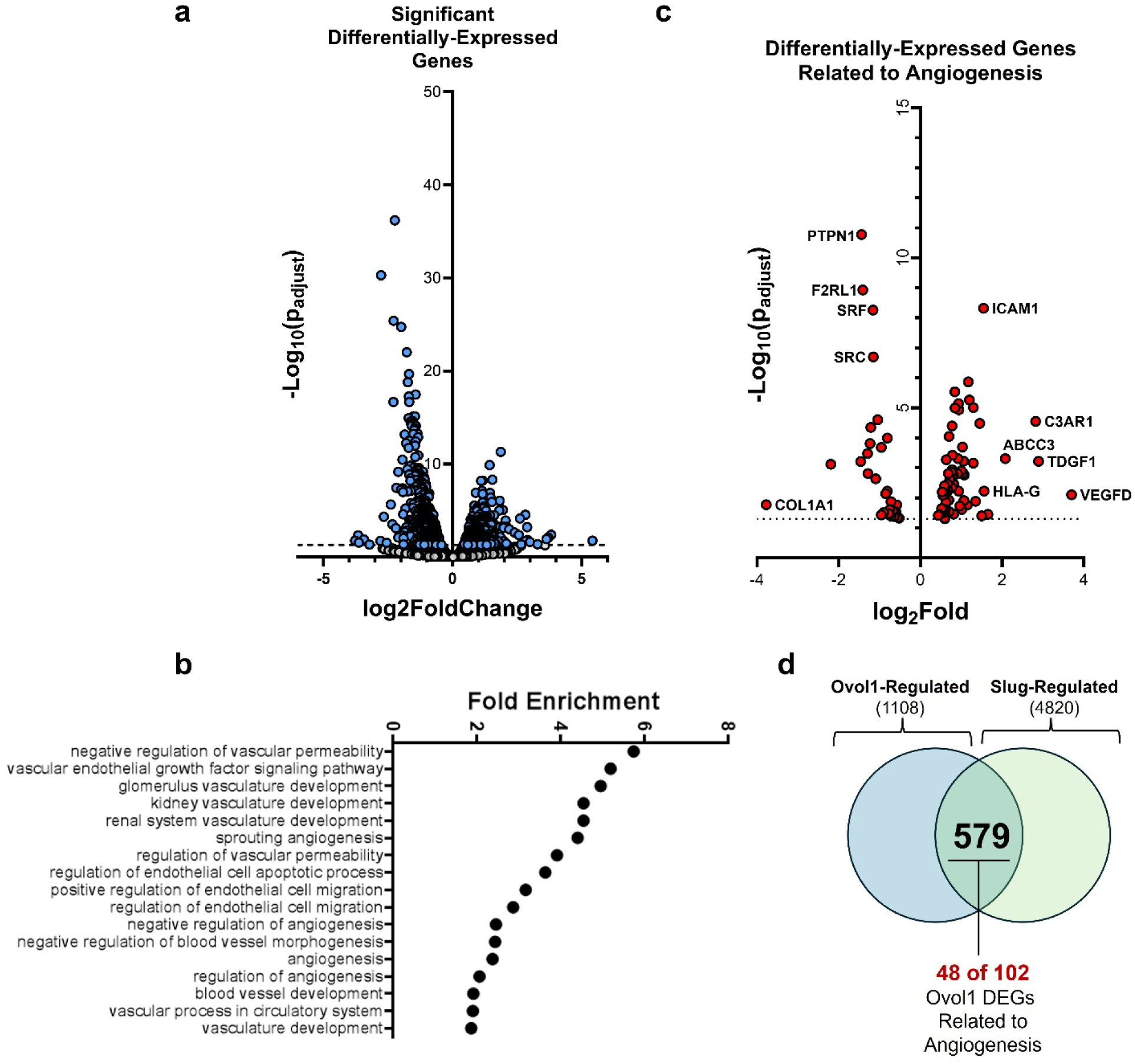
Ovol1 regulates genes associated with angiogenesis. To identify Ovol1-regulated genes and associated gene ontology pathways, we performed bulk transcriptomic analysis of HUVEC treated with si-Ovol1 to knockdown endogenous Ovol1. **a)** 1,108 significant (p<0.05) differentially-expressed genes (DEGs) were detected compared to si-Ctrl EC. **b)** Of the 518 total gene ontology (GO) terms identified as significantly enriched in these DEGs, 17 are specifically related to angiogenesis and vascular function. **c)** These angiogenesis-specific GO terms are driven by differential expression of 102 specific angiogenesis-associated genes. **d)** 579 (i.e., 52%) of the 1,108 total Ovol1-regulated genes identified in this dataset are also regulated by Slug (dataset published in Hultgren et al. 2020), including 48 (∼44%) of the 102 angiogenesis-associated genes from **b-c**.

We previously showed that the Slug transcription factor aregulates sprouting angiogenesis via activation of pEndoMT.[7,30] Comparison of that earlier RNASeq dataset identified 579 (i.e., 52%) of our Ovol1-regulated DEGs that overlapped with Slug-dependent genes we previously identified in EC [7] (**Figure 2d**), including 48 (i.e., ∼44%) of the angiogenesis-related genes identified in **Figure 2c** (and **Supplementary Table 3**). The strong overlap between Ovol1-regulated and Slug-regulated DEGs in these independent datasets argue that endothelial Ovol1 and Slug likely participate in the same signaling axis to influence sprouting angiogenesis.

### Ovol1 is co-expressed with Slug in sprouting angiogenesis

We next performed immunofluorescence in angiogenic sprouts generated using the Bead Assay, and we compared the spatial expression of Ovol1^+^ EC versus those that express Slug. We observed that Ovol1^+^ angiogenic EC nuclei were typically also Slug^+^ (**Figure 3a**, red arrowheads). Quantification of these data confirmed a positive correlation between Ovol1 and Slug expression in EC nuclei of angiogenic sprouts (**Figure 3b**).

**Figure 3:**
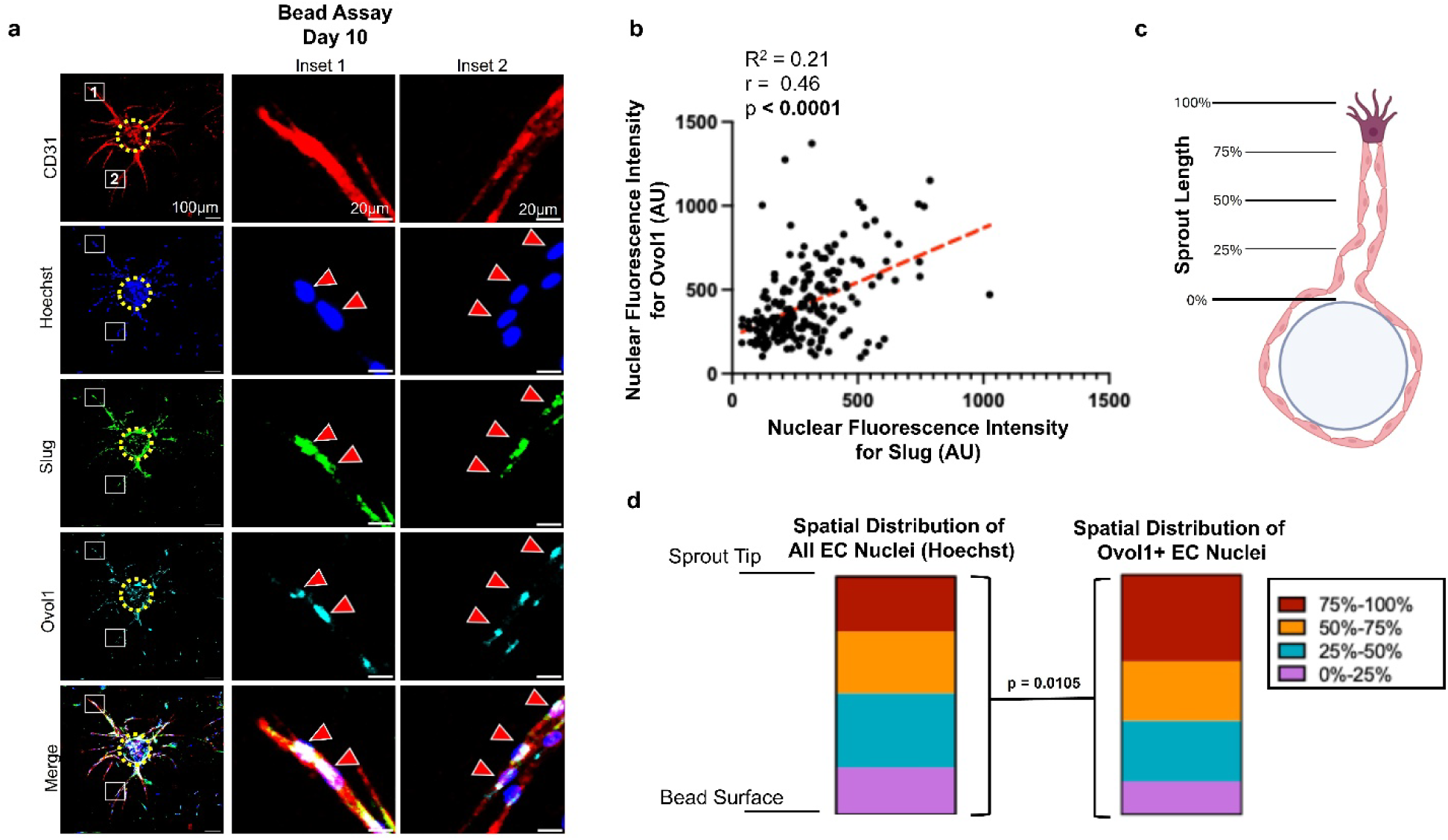
Ovol1 is co-expressed with Slug in angiogenic EC. **a)** Many angiogenic EC in Bead Assay (bead, yellow dashed circle) are positive for both (red arrowheads) nuclear Ovol1 (cyan) and nuclear Slug (green). **b)** Analysis of fluorescent intensity of individual angiogenic EC nuclei reveals a significant positive correlation between Ovol1 and Slug expression (pooled from sprouts from n=5 beads). To identify the spatial distribution of Ovol1^+^ EC within the sprout, **c)** we established quartiles relative to total sprout length. **d)** Compared to the distribution all EC nuclei across the sprout (which was relatively uniform), a significant difference was detected in spatial distribution of Ovol1^+^ EC, which appears to be driven primarily by enrichment of Ovol1^+^ EC in the distal sprout quartile (i.e., red, 75-100% of total sprout length), as well as lower than expected representation of Ovol1^+^ EC at the base of angiogenic sprouts (i.e., purple, 0-25% of total sprout length) (pooled from n=10 beads). (Statistics: Chi-Square Test).

To further understand the spatial distribution of Ovol1^+^ EC along the angiogenic sprout, we established quartiles for each sprout (**Figure 3c**). Using a Hoechst nuclear marker, we first confirmed the distribution of total endothelial cells across each sprout quartile. Next, we measured the distribution of Ovol1^+^ EC and found they were more prevalent in the distal quartile of the sprout (75-100%, red), where tip and stalk cells support sprout elongation. By contrast, relatively few Ovol1^+^ EC mapped to the base of the sprout (0-25%, purple) (**Figure 3d**).

### Ovol1 directly binds to the Slug gene to promote Slug expression

In **Figure 1**, we identified that endothelial cells predominantly express a variant of Ovol1 that migrates at ∼25kDa. One possible explanation for this apparent size difference is that endothelial Ovol1 might be truncated, including in a manner that might abolish one or more of the zinc-finger binding domains needed for it to bind to and regulate target genes. To therefore exclude this possibility, we used an overexpression construct in which the Ovol1 coding sequence is fused with a C-terminal V5 tag (i.e., Ovol1^OE^).[15] Transduction of endothelial cells with Ovol1^OE^ still produces a ∼25kDa Ovol1 band that can be detected with an anti-V5 antibody (**Figure 4a**). 293T cells also predominantly express the low molecular weight variant of Ovol1. Transduction of 293T cells with Ovol1^OE^ also produces V5-reactive bands corresponding to both low (∼25kDa) and high (∼30kDa) molecular weights (**Figure 4a**). These data not only indicate that the ∼25kDa Ovol1 variant is C-terminal intact, but also indicates that low-molecular weight Ovol1 can be produced by expressing full-length Ovol1 coding sequence. This suggests a post-transcriptional mechanism may generate the low-molecular weight Ovol1 variant that predominates in endothelial cells.

**Figure 4:**
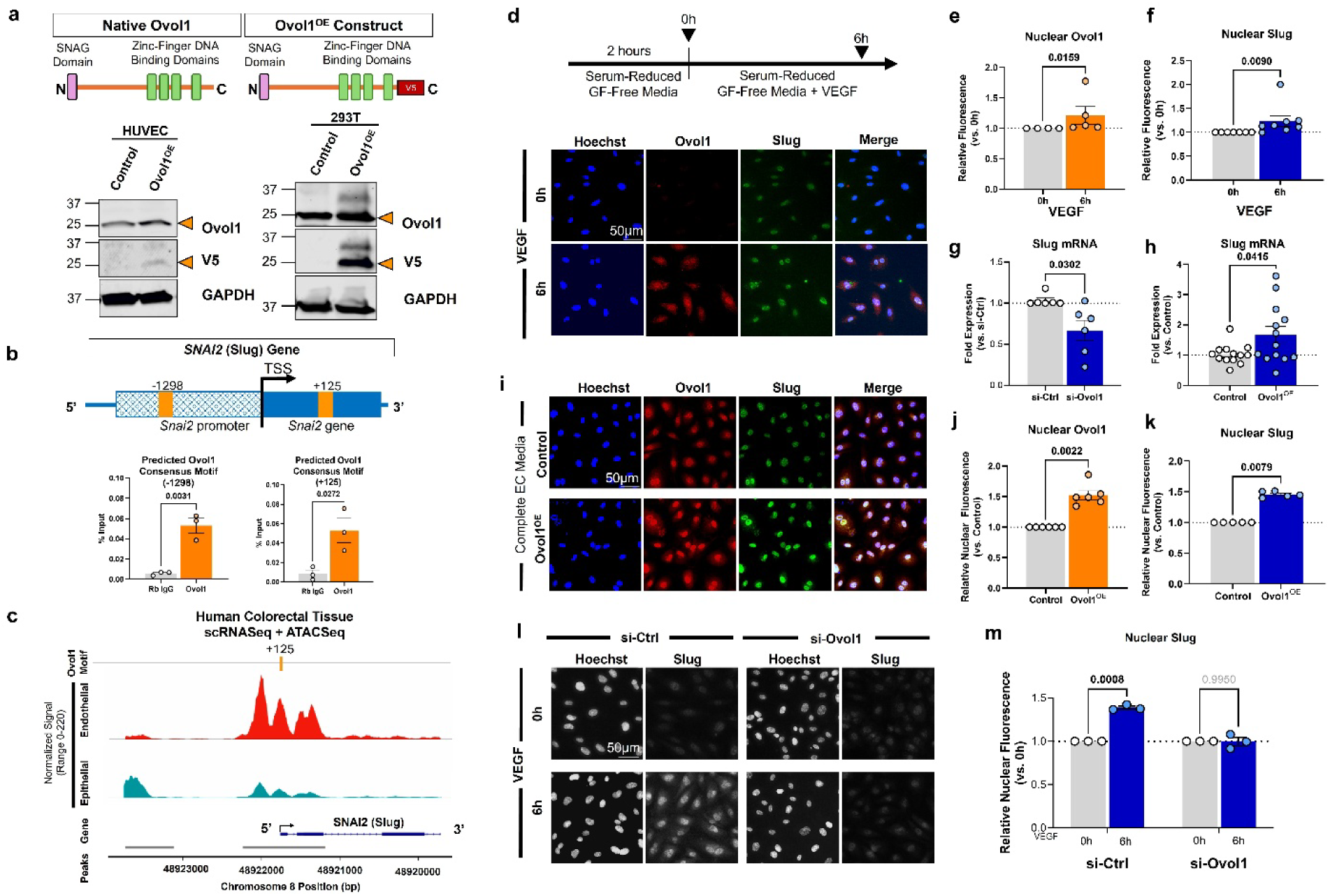
Ovol1 directly regulates Slug expression. **a)** Schematic of native Ovol1 reveals an N-terminal SNAG domain and several C-terminal zinc-finger binding domains. An Ovol1 overexpression construct contains the coding sequence for Ovol1 with a C-terminal V5 tag. Western blot of HUVEC transduced with the Ovol1^OE^ construct reveals V5-tagged Ovol1 predominantly running at ∼25kDa, whereas both 25kDa and 30kDa bands were detected in HEK293T Ovol1^OE^ cells. Blots are representative of 2-3 independent replicates. **b)** The Slug (Snai2) gene contains two predicted Ovol1-binding consensus sites flanking the transcriptional start site (TSS). ChiP-qPCR analysis reveals that Ovol1 is capable of binding to both consensus sites to pull down the Slug gene. **c)** In a single-nuclei multiomic sequencing[21] (GEO GSE29455) that captures both endothelial and epithelial cells (as well as other cell populations) from healthy human colon, our reanalysis identifies accessible chromatin at the Slug gene corresponding to the +125 Ovol1 consensus site and reveals higher single-cell ATACseq peak size for endothelial cells in healthy colon compared to adjacent colorectal epithelial cells. **d)** HUVEC monocultures were serum- and growth-factor starved for 2h, followed by treatment with 10ng/mL VEGF-A for 6h. Mean nuclear fluorescence for **e)** Ovol1 (n=4 independent replicates per group) and **f)** Slug (n=8 independent replicates per group) were both significantly increased at 6h VEGF (Statistics: Wilcoxon rank-sum test). **g)** Silencing of Ovol1 significantly decreased Slug mRNA in endothelial cells cultured in complete endothelial cell media (n=6 independent replicates per group, Statistics: Unpaired Students’ t-test), while **h)** Ovol1 overexpression significantly increased Slug mRNA compared to cells transduced with control lentivirus (n=13 independent replicates per group, Statistics: Wilcoxon rank-sum test). **i-k)** To confirm this observation at the protein level, we used immunofluorescence and observed increased nuclear expression of both Ovol1 (n=6 independent replicates per group, Statistics: Wilcoxon rank-sum test) and Slug (n=5 independent replicates per group, Statistics: Wilcoxon rank-sum test) in endothelial cells cultured in (growth factor-rich) complete EC media. **l-m)** Silencing of Ovol1 prevented VEGF-induced upregulation of nuclear Slug (n=3 independent replicates per group, Statistics: 2-way ANOVA with post-hoc t-test adjusted for multiple comparisons).

Since endothelial Ovol1 is C-terminal intact, we next explored whether endothelial Ovol1 directly regulates the Slug gene. Consensus sequence scanning of the *Snai2* gene (which encodes for Slug) identifies two high-probability Ovol DNA-binding motifs [i.e., C(C/T)G(A/T)TA][31,32] – one at position −1298 relative to the Slug transcriptional start site within the upstream promoter and enhancer region, and another at position +125 within the 5’ untranslated region of the Slug gene. To test if Ovol1 can functionally bind to either motif in the Slug gene, we performed ChIP-qPCR in endothelial cells using an anti-Ovol1 antibody and pulled down genomic DNA from both consensus motifs. Next, we performed sequence alignment and found that the +125 Ovol1 consensus motif was conserved in the Slug gene across mouse, rat, chimpanzee and human genomes (**Supplementary Figure 5a**), as well as broadly conserved across vertebrate species (**Supplementary Figure 5b**). To explore if the +125 Ovol1 binding motif is accessible in human cells, we reanalyzed a single-cell multiomic dataset generated from donor colorectal tissue, and which includes both endothelial and epithelial cell types. We performed PCA analysis to identify endothelial and epithelial cell populations based on marker expression, and compared ATAC-seq chromatin accessibility peaks for the Slug gene in these two cell types. We found that the +125 Ovol binding site of the Slug gene is significantly more accessible in endothelial cells compared to epithelial cells (**Figure 4c**).

Sprouting angiogenesis is regulated by soluble growth factors including vascular endothelial growth factor (VEGF).[3,33] We previously reported that VEGF upregulates Slug transcript levels in EC.[30] To test if Ovol1 is also involved in VEGF-induced upregulation of Slug, we cultured endothelial cells in serum-reduced and growth factor-free endothelial cell media for 2 hours prior to stimulation with exogenous VEGF. Interestingly, in the absence of serum and exogenous growth factors, baseline Ovol1 expression was not detectible (vs. **Figure 1b**) suggesting sensitivity of Ovol1 expression to these soluble signals. Stimulation with VEGF produced a nearly two-fold upregulation of nuclear Ovol1 within 1h of VEGF treatment (**Supplementary Figure 6a-b**) that began returning to baseline by 6h (**Figure 4d-e**). By contrast, Slug was more delayed in its upregulation following VEGF treatment: nuclear Slug was not significantly upregulated at 1h or 3h VEGF (**Supplementary Figure 6c**) and was only significantly increased at 6h VEGF (**Figure 4f**). Thus, VEGF upregulation of Ovol1 appears to precede upregulation of Slug, suggesting Ovol1 binding to the Slug gene might mediate this effect.

To test if Ovol1 is necessary and sufficient to upregulate Slug at the transcriptional level, we first used silencing RNA to knockdown endogenous Ovol1 in endothelial cells. This produced a significant decrease in Slug transcript levels (**Figure 4g**). Next, we used the Ovol1^OE^ lentiviral construct to constitutively express Ovol1, which produced a ∼50% increase in nuclear Ovol1 expression relative to control lentivirus (**Supplementary Figure 3c-f**). Ovol1 overexpression was sufficient to significantly upregulate Slug mRNA levels (**Figure 4h**). To confirm that this also resulted in a functional increase in nuclear Slug, we used immunofluorescence to measure Slug expression within the nuclei of Ovol1^OE^ endothelial cells and found both nuclear Slug (and nuclear Ovol1) were elevated with Ovol1 overexpression when compared to either untransduced EC (**Supplementary Figure 7**) or EC transduced with a control lentivirus (**Figure 4i-k**). Lastly, we investigated if Ovol1 is necessary for VEGF-induced upregulation of nuclear Slug. We used silencing RNA to prevent VEGF-induced Ovol1 upregulation (**Supplementary Figure 8**). We then compared nuclear Slug expression at 0h vs. 6h VEGF. Similar to what we observed for untransfected EC (**Figure 4d-f**), nuclear Slug was significantly upregulated in si-Ctrl EC at 6h VEGF, but this response was abolished in si-Ovol1 EC (**Figure 4l-m**). Taken together, our data indicate that Ovol1 is both necessary and sufficient for VEGF to upregulate Slug in endothelial cells.

### Ovol1 regulates sprouting angiogenesis via Slug in vitro

We previously showed that Slug is required for angiogenic sprouting in the *in vitro* sprouting angiogenesis Bead Assay,[30] whereas overexpression of Slug produces a hypersprouting phenotype (or complete mesenchymal transition) in a dose-dependent manner.[7] Based on this and our above observation that Ovol1 regulates VEGF-induced Slug (**Figure 4**), we next tested involvement of Ovol1 in angiogenesis. We again used silencing RNA to knockdown endogenous Ovol1. We found that angiogenic sprouting was significantly abrogated in si-Ovol1 EC by Day 6 in the Bead Assay (**Figure 5a**), characterized by significant reductions in both mean sprout number (**Figure 5b**) and sprout length (**Figure 5c**). On the other hand, EC transduced to constitutively overexpress Ovol1 resulted in a hypersprouting phenotype by Day 6 in the Bead Assay (**Figure 5d**), which was quantified as an increase in mean sprout number (**Figure 5e**) and sprout length (**Figure 5f**).

**Figure 5.**
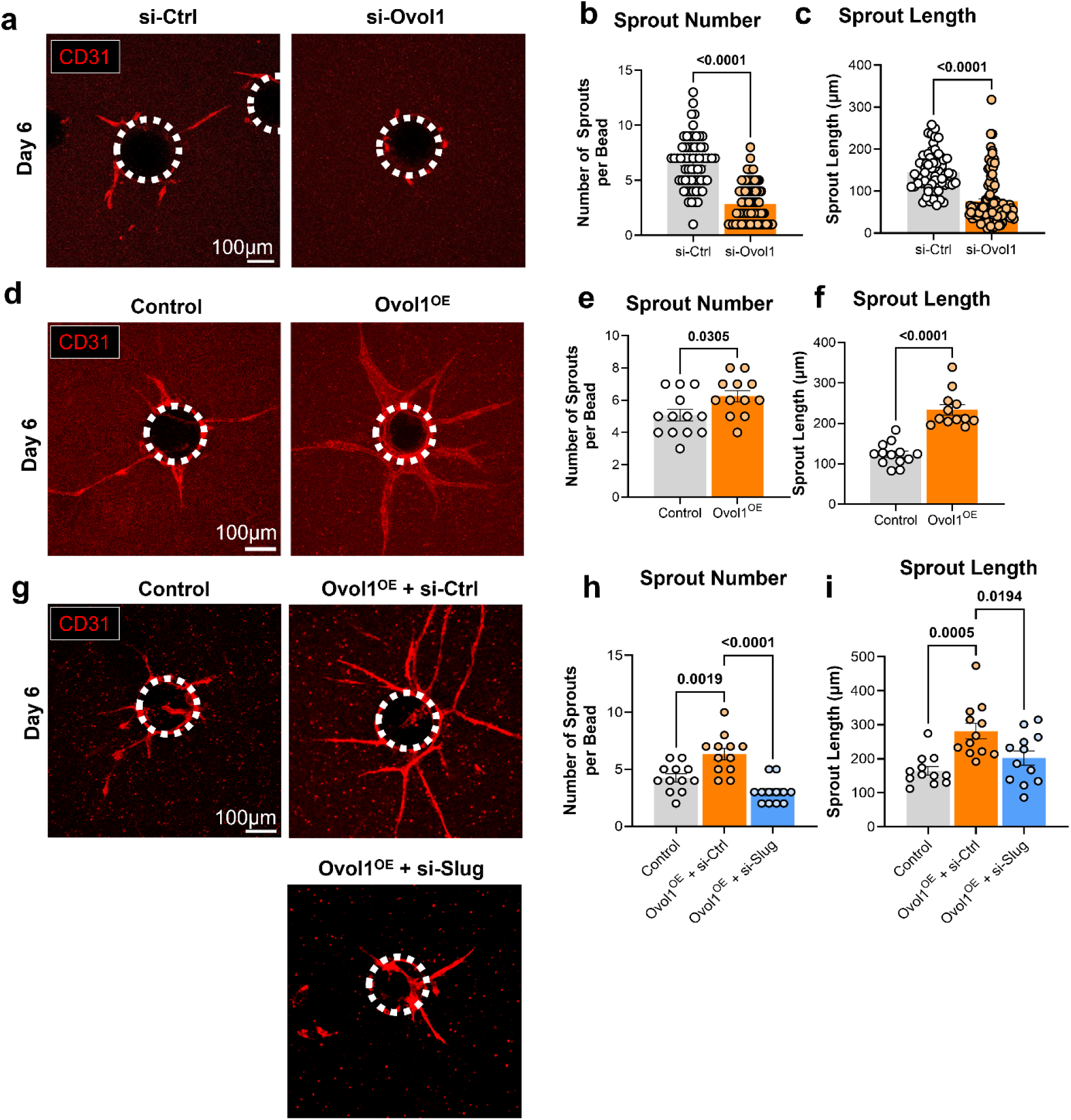
Ovol1 regulates angiogenesis via Slug. **a)** si-Ovol1 HUVEC fail to sprout in Bead Assay (bead, dashed circle) at Day 6 compared to si-Ctrl HUVEC, quantified as significant decreases in mean **b)** sprout number and **c)** sprout length (n=63 si-Ctrl, n=101 si-Ovol1 beads pooled across 9 independent replicates). (Statistics: Unpaired Students’ t-test) **d)** Ovol1^OE^ HUVEC undergo more aggressive sprouting compared to control (eGFP) HUVEC, quantified as significantly increased **e)** sprout number and **f)** sprout length (n=13 beads per group pooled across 3 independent replicates). (Statistics: Unpaired Students’ t-test). **g)** Silencing of endogenous Slug expression (via si-Slug) in Ovol1^OE^ HUVEC rescued both **h)** sprout number and **i)** sprout length compared to Ovol1^OE^cells transfected with a non-targeting siRNA control (si-Ctrl). (n=12 beads per group pooled across 3 independent replicates). (Statistics: 1-way ANOVA with post hoc t-test adjusted for multiple comparisons).

To test if Ovol1 regulates angiogenesis via Slug, we used silencing RNA to knockdown endogenous Slug (>85% efficiency, **Supplementary Figure 9**) in control or Ovol1^OE^ EC prior to seeding into the Bead Assay. We found that Slug knockdown normalized angiogenic sprouting of Ovol1^OE^ EC (**Figure 5h**). Sprout number and sprout length were not significantly different between Slug-deficient Ovol1^OE^ EC and control EC, whereas both were significantly reduced compared to Slug-intact Ovol1^OE^ EC (**Figure 5g-i**). Taken together, these data show that Ovol1 regulates angiogenesis *in vitro* via Slug.

We previously showed that Slug regulates angiogenesis by driving a marker profile consistent with pEndoMT – which is defined as the observed upregulation of some (but not all) mesenchymal markers alongside preservation of some (but not all) endothelial markers. [7] To test if Ovol1 also drives pEndoMT, we used qPCR to measure expression levels of classic mesenchymal and endothelial markers in control vs. Ovol1^OE^ EC. Ovol1 induced significant upregulation of mesenchymal markers N-Cadherin (*CDH2*), and Collagen Type VIII (*COL8A1*) and significant downregulation of Vimentin (*VIM*). Smooth muscle α-Actin (*ACTA2*) and Collagen Type I (*COL1A1*) were unaffected (**Figure 6a-e**). Meanwhile, Ovol1 overexpression preserved expression of classic endothelial markers CD31 (*PECAM1*) and VE-Cadherin (*CDH5*), while significantly reducing expression of VEGFR2 (*FLK1*) and Angiopoietin-1 (*ANGPT1*). A trend towards decreased Angiopoietin-2 (*ANGPT2*) was also observed, although this did not achieve statistical significance (**Figure 6f-j**).

**Figure 6.**
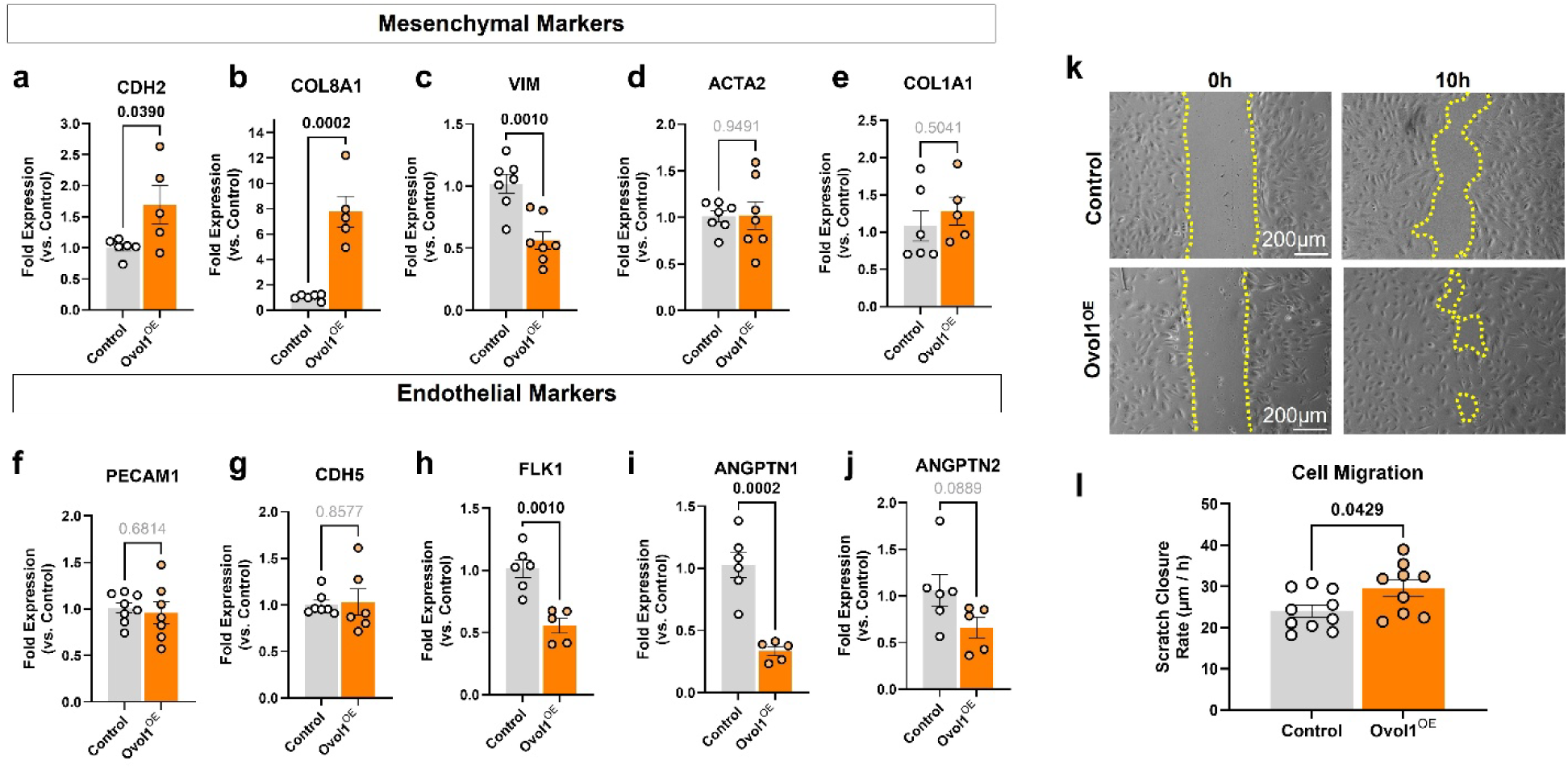
Ovol1 alters expression of endothelial and mesenchymal markers and promotes EC proliferation and migration. Mesenchymal transition markers **a)** CDH2 (N-Cadherin) and **b)** Collagen 8A (COL8A1) were significantly increased in Ovol1^OE^ EC while **c)** Vimentin (VIM) was significantly decreased vs. control EC. No significant differences were detected for **d)** smooth muscle α-actin (ACTA2) or **e)** Collagen 1A (COL1A1). In parallel, expression of EC markers **f)** CD31 (PECAM1) and **g)** VE-Cadherin (CDH5) were unaffected, while **h)** VEGFR2 (FLK1) was reduced. Vascular quiescence marker **i)** Angiopoietin 1 (ANGPTN1) was also significantly reduced while **j)** Angiopoietin 2 (ANGPTN2) also trended towards decreased expression. (n=5-7 independent samples per group, Statistics: Unpaired Students’ t-test). **k-l)** EC monolayer scratches were monitored for up to 10h, and scratch edges (dotted yellow lines) closed more rapidly for Ovol1^OE^ EC compared to Control EC. (n=10 Control, n=9 Ovol1^OE^ independent replicates per group; Statistics: Unpaired Students’ t-test).

Acquisition of a more mesenchymal-like pEndoMT endothelial cell state is expected to also promote more mesenchymal-like behaviors such as increased migratory capacity, a process critical for progression of sprouting angiogenesis.[3] Using a scratch assay approach, we quantified migration rate in confluent Ovol1^OE^ EC vs. control cell monolayers. We identified a significantly increased migratory capacity for Ovol1^OE^ EC (**Figure 6k-l**). Altogether, these data support the idea that Ovol1 promotes pEndoMT and increased cell migration.

### Endothelial Ovol1-Slug is largely dispensable for developmental angiogenesis

Given the striking requirement of Ovol1-Slug signaling for angiogenic sprouting in the Bead Assay (**Figure 5**), we next set out to test if this Ovol1-Slug axis is necessary for developmental angiogenesis in mice. We crossed Ovol1^flox/flox^ mice[22,23] with mice containing a conditional endothelial-specific (driven by *Cdh5* (VE-Cadherin) promoter) Cre driver[24] to generate the endothelial-specific Ovol1 knockout mouse line (i.e., Ovol1^ECKO^). We confirmed targeted endothelial-specific ablation of the Ovol1 gene by sorting lung endothelial cells (CD31^+^/CD45^−^ as previously described[25]) and performing endpoint PCR on isolated genomic DNA using the Ovol1 flanking primers in **Table 1** (**Supplementary Figures 10-11**). Endogenous Ovol1 amplified in Cre-negative animals but failed to generate a strong amplicon in both Cre-positive animals indicating successful recombination of the endogenous Ovol1 gene in sorted lung EC (**Supplementary Figure 11**). We further used immunofluorescence to compare frequency and distribution of Ovol1-positive EC nuclei in angiogenic and remodeling regions of the neonatal mouse retina between wild-type (Cre^−^) vs. Ovol1^ECKO^ (i.e., Cre^+^) littermates. Consistent with our earlier experiments, Ovol1^+^ EC nuclei represented ∼33% of all EC in the angiogenic front in Ovol1-intact (Cre-) mice with a >65% loss of Ovol1^+^ nuclei in Ovol1^ECKO^ littermates (**Supplementary Figure 11**). This confirms substantial penetrance of endothelial Ovol1 knockout in this line.

Given the severe phenotype of global Ovol1 knockout on overall tissue development,[12] we next set out to test if endothelial Ovol1 knockout affects overall animal viability. We confirmed that Ovol1^ECKO^ mice and wild-type (Cre-) littermates were generated at expected Mendelian ratios (**Supplementary Figure 12a**), that litter sizes were comparable between Ovol1^ECKO^ and wild-type C57/Bl6 breeding pairs (**Supplemental Figure 12b**), and that male-to-female pup ratio was approximately equal for Ovol1^ECKO^ litters (**Supplemental Figure 12c**). Together, these data indicate no significant embryonic loss for Ovol1^ECKO^ mice, consistent with the lack of obvious gross morphological defects for Ovol1^ECKO^ mice upon birth. We also compared animal weights between Ovol1^ECKO^ (vs. their wild-type littermates) in neonate and adult animals. Surprisingly, there was a noticeable (∼10%) increase in Ovol1^ECKO^ pup weight (vs. wild-type littermates) across both males and females (**Supplementary Figure 12d**), but this difference in animal weight was normalized by adulthood where genotype-specific differences were no longer detectible (**Supplementary Figure 12e**).

Using immunofluorescence and confocal microscopy, we compared the architecture of the developing neonatal retinal vasculature at post-natal day 6 between Ovol1^ECKO^ mice and wild-type (Cre-) littermates. We found that the retinal vasculature developed normally in Ovol1^ECKO^ mice (**Figure 7a**), with no discernible defects in vascular outgrowth (**Figure 7b**), vessel area (**Figure 7c**), or arteriogenesis (**Figure 7d**). Higher magnification imaging of the angiogenic front further revealed no change in tip cell number (**Figure 7e**).

**Figure 7.**
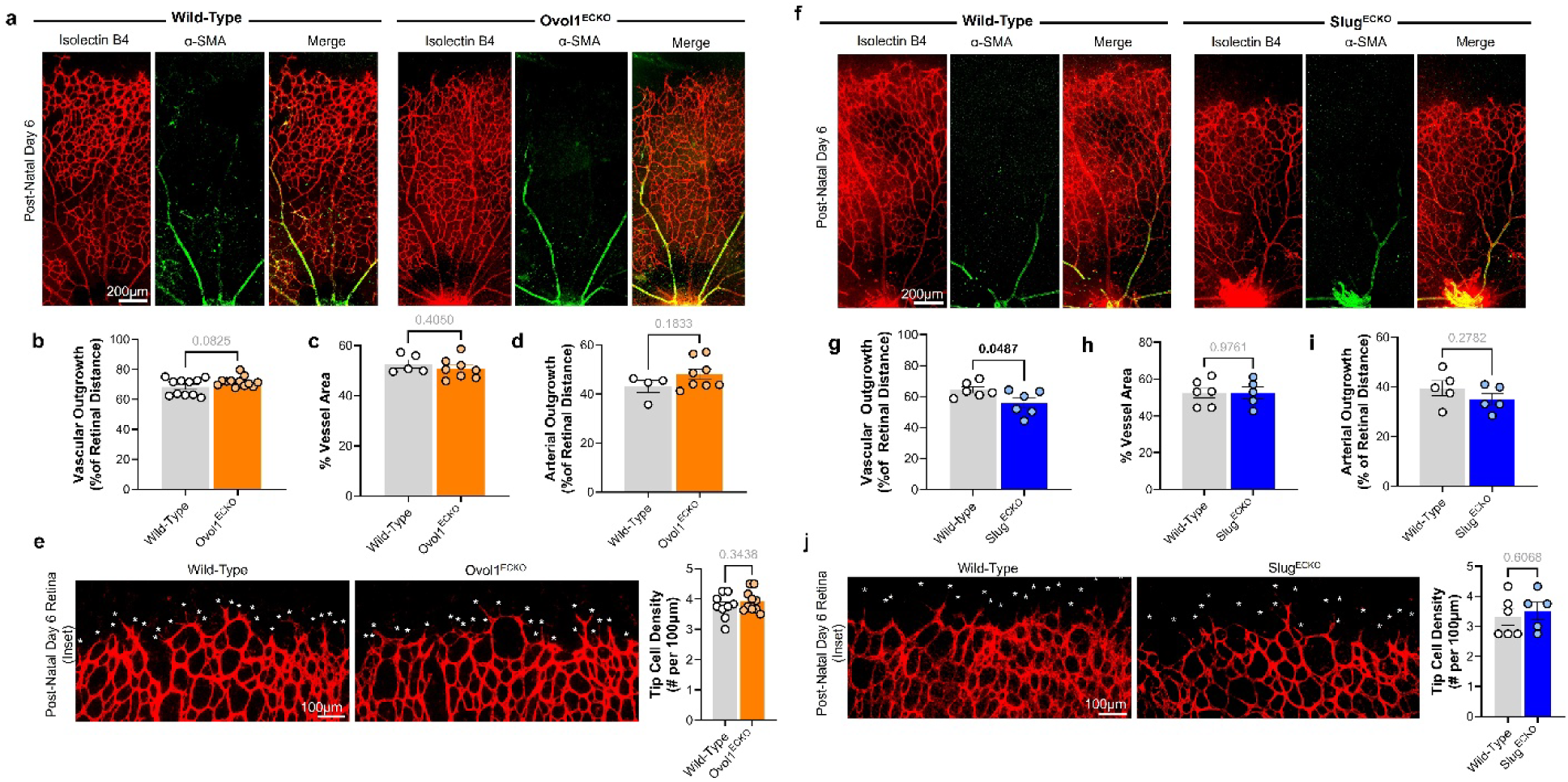
Endothelial Ovol1 is not required for developmental angiogenesis in the retina. **a)** Endothelial (isolectin B4, IB4) and vascular smooth muscle (α-SM-actin) staining of P6 retinas from Ovol1^ECKO^ and wild-type littermates reveal no significant differences in **b)** vascular outgrowth (n=12 wild-type, n=11 Ovol1^ECKO^ mouse littermates, Statistics: Unpaired Students’ t-test), **c)** vessel area (n=5 wild-type, n=8 Ovol1^ECKO^ mouse littermates, Statistics: Unpaired Students’ t-test), **d)** arterial outgrowth (n=4 wild-type, n=8 Ovol1^ECKO^ mouse littermates, Statistics: Unpaired Students’ t-test), or **e)** tip cell density (n=9, n=10 Ovol1^ECKO^ mouse littermates, Statistics: Unpaired Students’ t-test). **f)** In the P6 retina of Slug^ECKO^ mice, we observed **g)** significantly reduced vascular outgrowth (n=6 wild-type, n=6 Slug^ECKO^ mouse littermates) with no significant changes in **h)** vessel area (n=6 wild-type, n=5 Slug^ECKO^ mouse littermates), **i)** arterial outgrowth (n=5 wild-type, n=5 Slug^ECKO^ mouse littermates, Statistics: Unpaired Students’ t-test), or **j)** tip cell density (n=5 wild-type, n=5 Slug^ECKO^ mouse littermates, Statistics: Unpaired Students’ t-test).

We next asked if the lack of a developmental angiogenesis phenotype in Ovol1^ECKO^ phenocopies loss of endothelial Slug. We previously reported that global Slug knockout (Slug^KO^) mice – i.e., mice with germline knockout of the endogenous *Snai2* (Slug) gene – exhibit modest delays in developmental angiogenesis in the retina that resolve by later developmental timepoints.[7] To assess the precise contribution of endothelial Slug to this phenotype, we generated a novel Slug^flox/flox^ mouse line in which flanking loxP sites were inserted by CRISPR/Cas9 gene editing around the endogenous *Snai2* (Slug) gene locus. We crossed Slug^flox/flox^ mice with the same *Cdh5* promoter-driven Cre mouse as was used to generate the Ovol1^ECKO^ mouse line above. Similar to our observation in Ovol1^ECKO^ mice, amplification of the Slug gene from genomic DNA isolated from FACS-sorted lung EC (**Supplementary Figure 13**) was less robust in Slug^ECKO^ mice compared to (Cre-) wild-type littermate controls, although a faint amplicon could still be detected (**Supplementary Figure 14a-b**). In the angiogenic front of the P6 mouse retina, frequency of Slug^+^ EC was also reduced by ∼70% in Slug^ECKO^ mice compared to wild-type (Cre-) control littermates (**Supplementary Figure 14c**).

Similar to Ovol1^ECKO^, Slug^ECKO^ mice demonstrated no obvious gross defects. Slug^ECKO^ mice and wild-type (Cre-) littermates were generated at expected Mendelian ratios (**Supplementary Figure 15a**), and we observed no differences in average litter size (**Supplementary Figure 15b**) or male-to-female birth ratio (**Supplementary Figure 15c**) for Slug^ECKO^ litters compared to wild-type C57Bl/6 mice. Animal weight of Slug^ECKO^ mice was comparable to wild-type (Cre-) littermates for either neonates or adult mice (**Supplementary Figure 15d-e**). Lastly, similar to Ovol1^ECKO^ and our previous reports using global Slug^KO^ mice,[7] we observed no major failure of developmental angiogenesis in the P6 retina (**Figure 6f**). Consistent with the phenotype of Slug^KO^ mice, Slug^ECKO^ mice exhibited delayed vascular outgrowth (**Figure 7g**), but no effect on vessel area or arterial outgrowth (**Figure 7h-i**). Whereas we previously observed an increase in tip cell density in Slug^KO^ mice,[7] no differences in tip cell density were observed in Slug^ECKO^ animals (**Figure 7j**).

Taken together, these data show that Slug^ECKO^ mice phenocopy the delayed vascular outgrowth phenotype first reported in global Slug^KO^ mice,[7] but that developmental angiogenesis otherwise proceeds largely normally. Meanwhile, endothelial Ovol1 is completely dispensable for developmental angiogenesis in the retina.

### Endothelial Ovol1-Slug is essential for pathological angiogenesis

We previously showed that endothelial Slug is highly expressed in developing blood vessels. However, in the post-natal animal, healthy EC lack strong Slug expression, whereas Slug expression is reacquired in cancer where many tumor EC are Slug-positive. [7,30] Here, we report a similar finding for Ovol1. While Ovol1 was only weakly expressed in vascular endothelium of healthy mouse colon, endothelial Ovol1 expression was significantly increased in nuclei of tumor vessels that form in colorectal cancer syngraft tissue (**Figure 8a-c**). As expected, Ovol1 was also abundant in the surrounding healthy and cancerous epithelial cells. Therefore, to specifically visualize endothelial Ovol1 in tumors, we used a guinea pig ► mouse tumor xenograft model, in which host mouse vessels invade tumors grown from engrafted guinea pig cells. Resulting tumor tissue was probed using a mouse-specific Ovol1 antibody. Using this approach, we confirmed that Ovol1 expression clearly co-localized with CD31-positive tumor vessels (**Figure 8d-e**).

**Figure 8.**
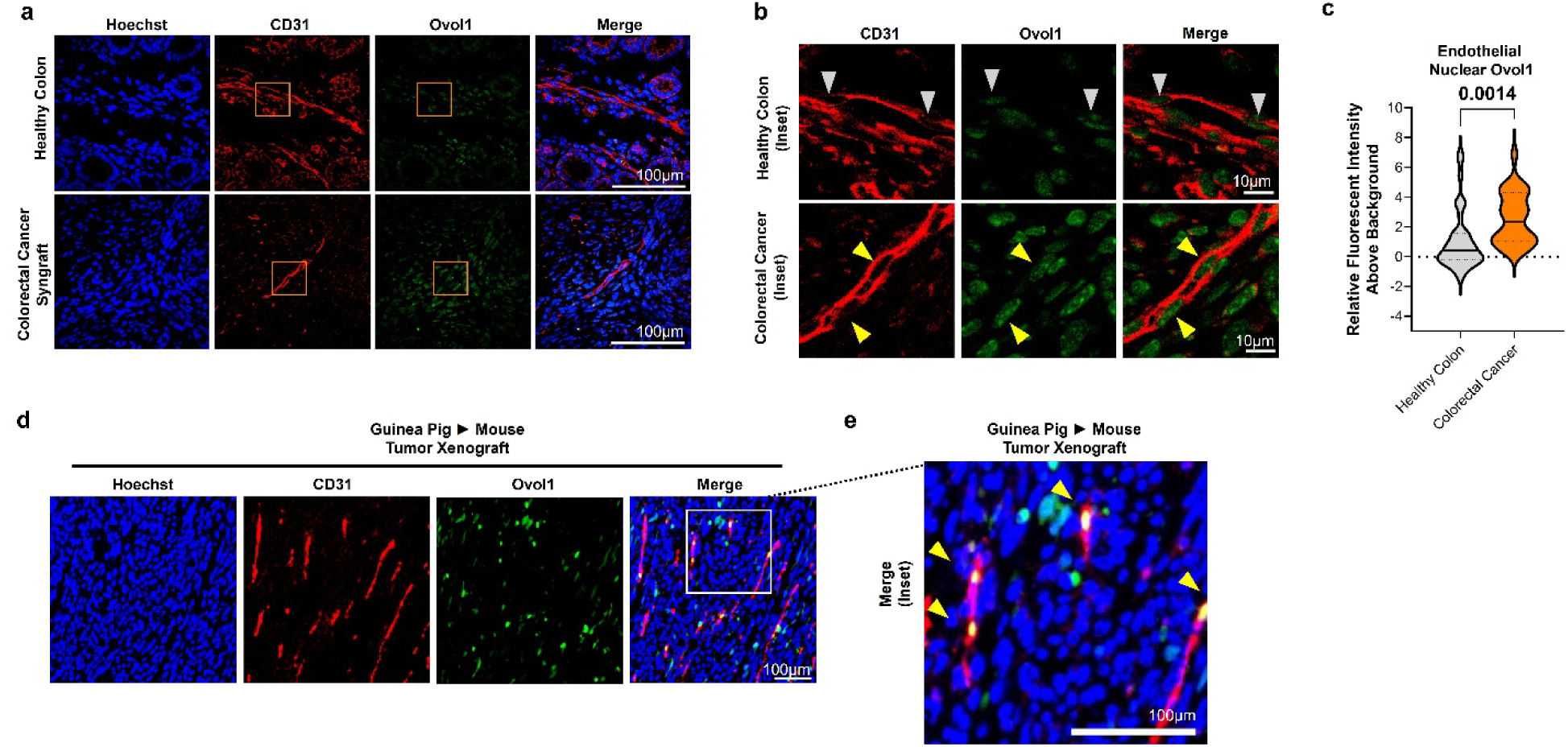
Endothelial Ovol1 is upregulated in tumor EC. **a-b)** Sections of colorectal cancer tumor syngraft and healthy colon from the same mouse were stained for Ovol1 (green) and endothelial marker CD31 (red). Nuclei (Hoechst, blue) of CD31^+^ vessels (inset, grey or yellow arrowheads) were individually examined for expression of nuclear Ovol1. **c)** Fluorescent intensity of nuclear Ovol1 was significantly increased in EC of CD31^+^ tumor vessels (n=39 cells) vs. EC of CD31^+^ vessels of healthy colon (n=27 cells). (Statistics: Unpaired Students’ t-test). To further visualize EC-specific Ovol1, we used a guinea pig tumor xenograft model, in which tumor vessels (but not stroma) would arise from mouse host tissue. **d-e)** In guinea pig ► mouse tumor xenograft, mouse Ovol1 expression co-localized primarily with CD31^+^ tumor vessels (yellow arrowheads).

To test if endothelial Ovol1 is required for tumor neovascularization, we repeated the tumor syngraft experiment in Ovol1^ECKO^ mice. Compared to tumors implanted into wild-type (Cre-) littermates, tumors failed to grow in Ovol1^ECKO^ mice (**Figure 9a**). This was not due to differences in animal weight as engrafted tumors grew (**Supplementary Figure 16**), indicating this was not due to a failure of experimental animals to thrive. Following harvest at 6 weeks, colorectal tumor syngrafts engrafted in Ovol1^ECKO^ mice generally appeared smaller and less well-perfused by blood (**Figure 9b**). Using immunofluorescent staining for the endothelial marker CD31 to visualize tumor vasculature, we found a significant reduction in tumor microvessel density in Ovol1^ECKO^ mice compared to tumors engrafted in wild-type control littermates (**Figure 9c-d**), associated with a significant decrease in tumor microvascular density. Similar results were observed for tumors engrafted in Slug^ECKO^ mice (**Figure 9e-h**). Importantly, failure of tumor syngrafts to grow could not be explained by a difference in circulating immune cell response: hematological profile was indistinguishable between Slug^ECKO^ mice vs. (Cre-) wild-type littermates (**Supplementary Figure 17**). Tumor syngrafts of a different murine colorectal cancer line (MC38) also failed to grow in Ovol1^ECKO^ mice (**Supplementary Figure 18**), indicating that these results were not cancer line dependent.

**Figure 9.**
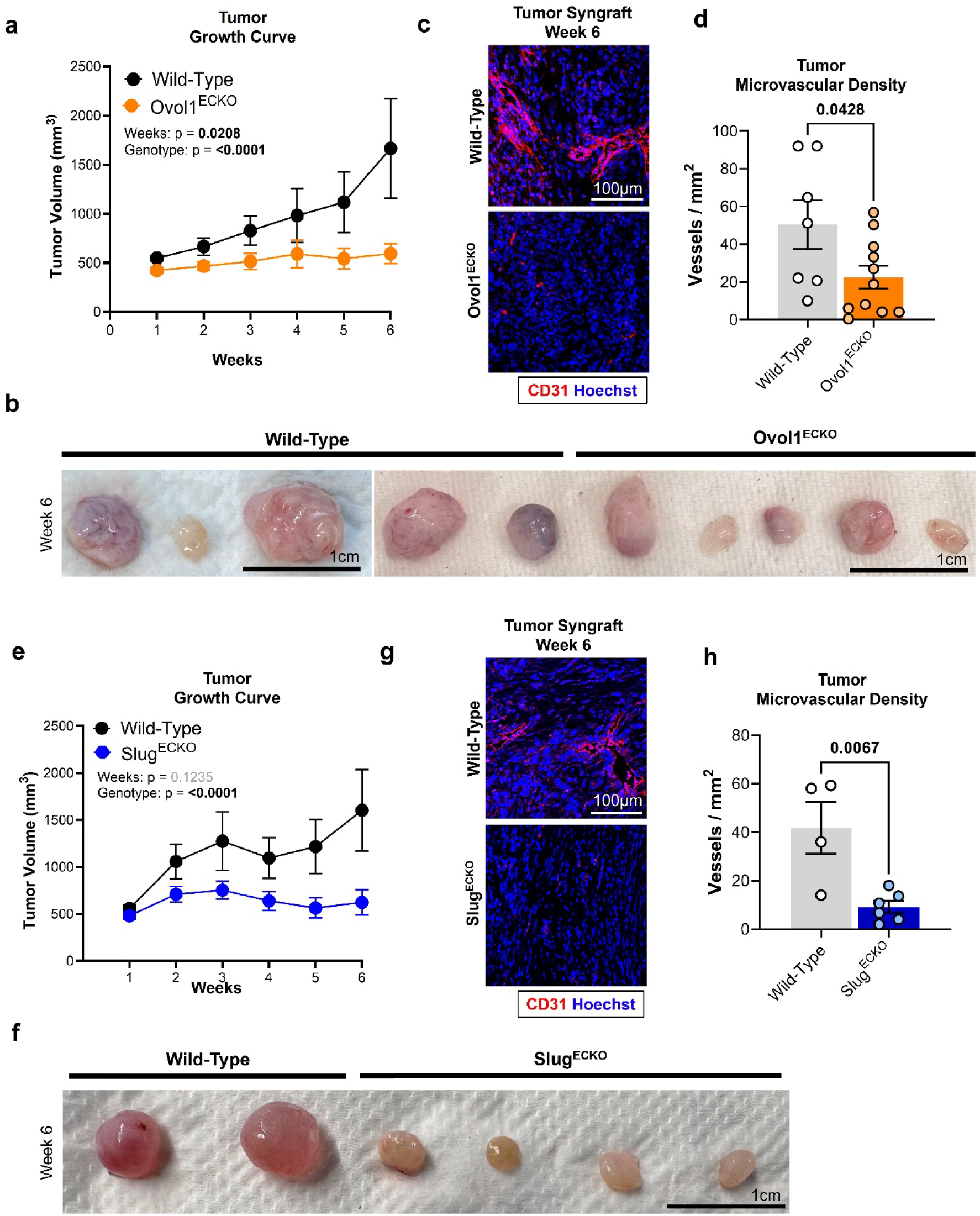
Endothelial Ovol and Slug are essential for pathological angiogenesis in cancer. **a)** CMT93 (colorectal cancer) tumor syngrafts injected into Ovol1^ECKO^ failed to grow compared to those injected into wild-type (Cre-) littermate controls (n=19 wild-type, n=17 Ovol1^ECKO^ mouse littermates), identified as a statistically significant effect of genotype in these studies. (Statistics: 2-way ANOVA). **b)** Tumors engrafted in Ovol1^ECKO^ mice appeared generally smaller than those engrafted in wild-type littermates. **c)** Immunofluorescent staining of week 6 syngraft tissue for CD31 (endothelial marker) identified fewer blood vessels in tumors from Ovol1^ECKO^ mice, corresponding with **d)** a significant decrease in tumor microvascular density (n=7 wild-type, n=10 Ovol1^ECKO^ mouse littermates). (Statistics: Unpaired Students’ t-test). **e)** CMT93 colorectal cancer syngrafts injected into Slug^ECKO^ mice also grew poorly over 6 weeks (n=19 wild-type, n=17 Slug^ECKO^ mouse littermates), identified as a statistically significant effect of genotype in these studies. (Statistics: 2-way ANOVA) and **f)** appeared smaller than those engrafted into wild-type (Cre-) littermate control animals. **g)** CD31-positive microvasculature was sparse in tumors engrafted in Slug^ECKO^ mice, corresponding with a **h)** significant decrease in tumor microvascular density (n=4 wild-type, n=6 Slug^ECKO^, Statistics: Unpaired Students’ t-test).

Lastly, to test requirement of the Ovol1-Slug pEndoMT signaling axis for pathological angiogenesis beyond the context of cancer, we used the oxygen-induced retinopathy (OIR) model, a widely-used model of pathological neovascularization in the eye (**Figure 10a**). Following exposure to hyperoxic conditions to induced vaso-obliteration followed by normoxia to stimulate neovascularization leading to closure of the avascular area (**Figure 10a**), we performed immunofluorescent staining of P17 retinas to visualize retinal vasculature (**Figure 10b**). We observed a significant increase in post-OIR avascular area at P17 for Slug^ECKO^ mice compared to wild-type (**Figure 10d**), indicating that Slug is also required for pathological angiogenesis in this widely-used model of hypoxic neovascular retinopathy.

**Figure 10.**
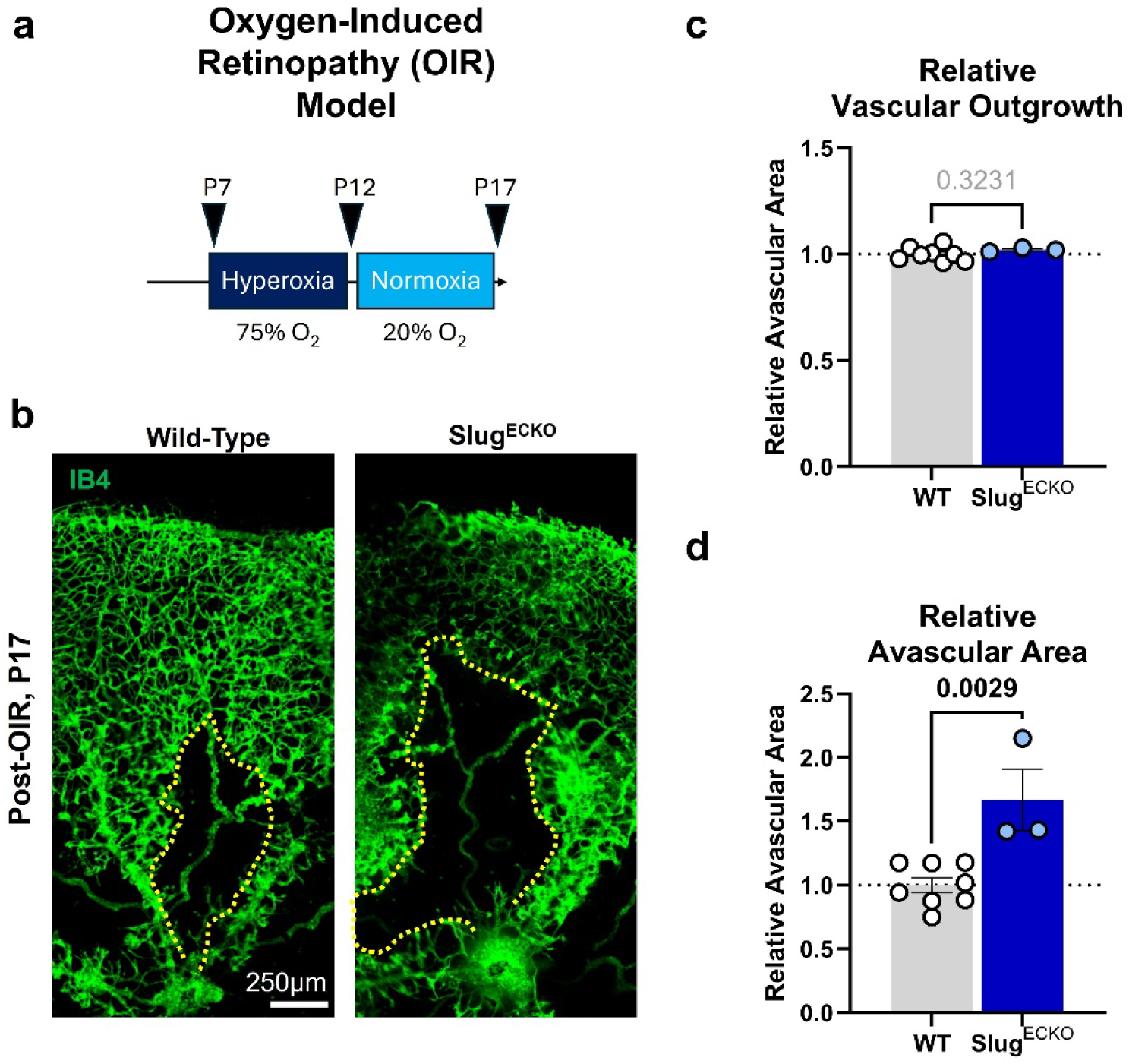
Endothelial Ovol and Slug are essential for neovascularization in the oxygen-induced retinopathy (OIR) model. **a)** In the oxygen-induced retinopathy (OIR) model of pathological neovascularization, P7 mice are placed with their dams in 75% O_2_ hyperoxic conditions for 5 days to induce vasoobliteration, followed by return to normal room air for an additional 5 days to stimulate neovascularization. **b)** Immunofluorescent staining of P17 retinas after OIR. **c)** Total vascular outgrowth is similar between wild-type and Slug^ECKO^ mice at P17 post-OIR (n=8 wild-type, n=3 Slug ^ECKO^ mouse littermates, Statistics: Unpaired Students’ t-test) but **d)** relative avascular area was significantly larger in Slug^ECKO^ compared to wild-type (Cre-) control littermate, indicating reduced pathological angiogenesis in Slug^ECKO^ mice (n=8 wild-type, n=3 Slug^ECKO^ mouse littermates, Statistics: Unpaired Students’ t-test).

## Discussion

In this study, we find that endothelial cells express a unique lower molecular-weight variant of Ovol1 that surprisingly promotes (rather than suppresses) transcription of target gene Slug and pEndoMT. Pro-angiogenic VEGF rapidly upregulates endothelial Ovol1 and downstream Slug to activate pEndoMT. Yet, while Ovol1-Slug signaling appears to be largely redundant for developmental angiogenesis and the Ovol1-Slug activation is absent from quiescent and established vessels of healthy tissue, we report that Ovol1 becomes reactivated in tumor vessels and that the endothelial Ovol1-Slug axis is almost completely essential for sprouting angiogenesis in cancer. Lastly, we also find that Slug is required for pathological angiogenesis in a model of hypoxic neovascularization in the retina.

During tissue development and in cancer, epithelial Ovol1 is activated by TGF-β[34] and Wnt[35] signaling to maintain cell commitment and critically suppress mesenchymal transition;[13] but, Ovol1’s involvement in endothelial cell types has not been well-characterized. In our studies, we find that endothelial cells express a distinct lower molecular-weight variant of Ovol1, and that Ovol1 in endothelial cells surprisingly promotes (rather than suppresses) transcription of target gene Slug and pEndoMT. Pro-angiogenic VEGF rapidly upregulates endothelial Ovol1 and downstream Slug to activate pEndoMT. Yet, while Ovol1-Slug signaling appears to be largely redundant for developmental angiogenesis and the Ovol1-Slug activation is absent from quiescent and established vessels of healthy tissue, Ovol1 becomes reactivated in tumor vessels and the endothelial Ovol1-Slug axis is almost completely essential for sprouting angiogenesis in cancer. Endothelial Slug is also required for pathological angiogenesis outside of the context of cancer, and neovascularization into the avascular area generated in the oxygen-induced retinopathy model is significantly reduced in endothelial-specific Slug knockout mice.

Our results have several major implications. First, they identify an Ovol1 variant that predominates in endothelial cells, and that appears to exert an unexpectedly opposite effect on Slug and pEndoMT vs. what would be expected based on Ovol1’s pro-epithelial role in epithelial cells.[14,9,13,10] Indeed, we initially hypothesized that – similar to in epithelial cancer cells – Ovol1 would suppress Slug and limit pEndoMT in endothelial cells. In support of this initial idea, Chen and colleagues recently identified Slug as one of several direct Ovol1 target genes in keratinocytes (an epithelial cell type).[14] Further work by Roca and colleagues shows that in epithelial cancer cells Ovol1 suppresses (not activates) Slug transcript with no appreciable effect on Slug protein in epithelial cells.[10] This may occur through active recruitment of HDAC1 by Ovol1’s N-terminal SNAG domain (peptide sequence: MPASFLVK[36]) and/or by passive competition for a common DNA consensus motif (core nucleotide sequence: CC/TGTTA[32]).

Although Ovol1 is an anti-mesenchymal transcriptional repressor in epithelial cells, we were surprised to that in *endothelial* cells, Ovol1 promotes – rather than suppresses – Slug expression and upregulates mesenchymal-like marker expression and pro-migratory cell behavior. How Ovol1 might produce such distinct outcomes in epithelial vs. endothelial cell types is still unclear, but our data implicate a distinct function of the (∼25kDa) low-molecular weight variant of Ovol1 that predominates in endothelial cells. We show that the ∼25kDa Ovol1 variant of endothelial cells can be produced from full-length Ovol1 coding sequence, is C-terminal intact, and retains functional DNA-binding capacity to the Slug gene. These data together suggest a post-transcriptional mechanism of generating the ∼25kDa Ovol1 variant from full-length mRNA. One intriguing possibility is that the ∼25kDa Ovol1 variant arises through an alternative internal translational start site that is favored in endothelial cell types, but perhaps not in epithelial cancer cell types. This idea is supported for other proteins in other tissues: for example, recent work by Lee and colleagues reveal how alternative ribosomal translational start sites in the synaptic protein neuronal pentraxin receptor (NPR) produce diverse protein variants (i.e., proteoforms) with differing expression patterns and functions in the brain.[37] Full-length Ovol1 mRNA (Genbank NM_004561.4) contains two translational start sites – i) the primary translational start site (i.e., Kozak sequence) at position 312, and ii) an internal predicted translational start site at position 498 (independently predicted by both NetStart and ATGpr tools). Initiation of translation at the latter start site would produce an N-terminal truncated variant of Ovol1 (Genbank XP_05422959) with an amino acid length of 205 residues (vs 267 in full-length Ovol1) and a predicted molecular weight of 23.32 kDa (ExPasy). Strikingly, this is consistent with the apparent molecular weight of the Ovol1 variant we observe in endothelial cells (**Figure 1a**). N-terminal truncation of Ovol1 would abolish the SNAG domain while preserving intermembrane and C-terminal zinc-finger DNA-binding domains (**Supplementary Figure 19a-d**). In epithelial cells, the Ovol1 SNAG domain This may occur through active recruitment of HDAC1 by Ovol1’s N-terminal SNAG domain (peptide sequence: MPASFLVK[36]) is essential for recruiting transcriptional co-repressors and suppressing target gene transcription in epithelial cells[32,14] (**Supplementary Figure 19a-b**); loss of this domain might produce alternative outcomes on target gene transcription. The mechanism by which an N-terminal truncated Ovol1 variant might be preferentially expressed in some cell types (e.g., endothelial cells and others) but not others (e.g., epithelial cancer cells) is still unclear. Possibilities include transcript variants that emerge from alternative splicing or alternative transcriptional start sites, and/or alternative translation through the binding of RNA-binding proteins. On the other hand, our data might also be explained by differences in Ovol1 phosphorylation leading to altered protein migration. Indeed, Ovol1 contains several phosphorylatable serine/threonine residues, and Vetere and colleagues speculated that distinct banding patterns for Ovol1 in Western blots from different epithelial tissue types in rat may represent tissue-specific expression of Ovol1 in distinct phospho-states.[29] Additional experiments are clearly required to distinguish between these various possibilities.

The discrepancy in Ovol1 effects on Slug expression between endothelial vs. epithelial cells may also be related to differences in Ovol1 binding motif accessibility in the Slug gene between these two cell types. Indeed, our reanalysis of single-cell multomics data shows that the Ovol1 binding site on the Slug gene is more accessible in endothelial cells rather than epithelial cells. We also find that endothelial Ovol1-Slug signaling is critical in pathological angiogenesis in both cancer and retinopathy models of pathological angiogenesis, whereas both transcription factors are largely dispensable for developmental angiogenesis. While the lack of a severe developmental phenotype in our endothelial-specific knockout animals may be in part explained by the incomplete penetrance of Cre recombination in these animals, this cannot fully explain why pathological angiogenesis in tumor syngraft and OIR models are so profoundly reduced in our knockout animals. Instead, our data are consistent with the idea that the Ovol1-Slug axis is redundant or dispensable during developmental angiogenesis (perhaps due to compensatory pro-angiogenic pathways) but becomes distinctly essential in neovascular disease - likely as a consequence of its pathological reactivation to centrally control pro-mesenchymal signaling. Recent sequencing by Yang and colleagues reveals that a significant proportion of tumor endothelial cells acquire a mesenchymal-like gene signature and identity,[8] and in both this work and our earlier work, as well as the work of others, it is revealed that master epithelial-to-mesenchymal transition transcription factors – such as Slug, Snail, and Twist – that drive mesenchymal transition in epithelial cells are also essential for establishment of tumor blood vessels, where Snail has also been shown to confer resistance to anti-angiogenic drugs.[38,8,7,2]

In epithelial cells, epithelial-to-mesenchymal transition (EMT) is governed by a dense, interconnected network of these EMT master transcription factors (which notably includes both Slug and Ovol1) and microRNAs all of which are regulated at the epigenetic, transcriptional, post-transcriptional, and post-translational levels to coordinate cell state transitions. The presence of multiple mutual inhibition and positive feedback motifs (e.g., as described by Zhang and colleagues in epithelial cell types [39]) within this interaction network can give rise to multiple stable states[40] – including bistable hybrid states – that can simultaneously emerge in response to the same signaling conditions based on an individual cell’s regulatory protein levels,[41] and/or its microenvironmental and epigenetic context cues. In disease, aberrant activation of the pEndoMT regulatory network – such as aberrant upregulation of Ovol1 and/or miR-150 also recently shown to regulate Slug [42] – might lead to the emergence of disease-specific bistable states that drive pathological neovascularization. Alternatively, key target genes of Ovol1 may have multiple functional enhancer elements (such as shadow promotors[43]) that are operational during development, and Ovol1-independent ones may become inaccessible post-developmentally. Further studies are needed to understand precisely how the healthy vs. diseased microenvironment alters requirement for Ovol1-Slug signaling in endothelial cells.

In conclusion, we report a novel Ovol1-Slug signaling axis that regulates sprouting angiogenesis *in vitro* and tumor neovascularization *in vivo*. These data suggest that the functional requirement for endothelial Ovol1-Slug may be specific to pathological angiogenesis. This raises the exciting possibility that the Ovol1-Slug axis (and/or its downstream genes) could be leveraged to selectively ablate pathological neovascularization in several diseases – including cancer – while potentially leaving healthy vasculature intact.

## Acknowledgements

The authors would like to acknowledge Dr. Nan Wu Hultgren (UCLA) for her feedback on this work, and Emma Bates (Tulane) for technical support in some studies. The authors would like to acknowledge the Transgenic Mouse Facility at UCI for their help in developing the Slug^flox^ mouse, as well as technical support from Dr. Jovanny Zabaleta and Dr. Jone Garai of the Translational Genomics Core at the Louisiana Cancer Research Center (LCRC), and Constance P. Porretta of the Tulane Flow Cytometry and Cell Sorting Facility. YCCEL1 cells were a gift of E. Flemington (Tulane). Lastly, the authors would like to acknowledge animal care technicians at Tulane, UCI, and LSU for their care of our research animals, as well all other members of the Fang lab for their invaluable scientific discussion and feedback on this project. Certain explanatory cartoons were generated using Biorender.

## Author Contributions

Data were collected and analyzed by KT, BB, HC, SS, MP and JSF. KT and JSF wrote the manuscript. BAS performed guinea pig tumor xenograft studies, RKC and SMM contributed to RNA sequencing sample collection, and CJH and MCB helped in next generation sequencing analysis. NC, AK, and KS performed sequence alignments and protein structure predictions. DPH, SMM, and CCWH provided scientific input and manuscript revisions. JSF, BAS, DPH, and CCWH secured funding for this work.

## Data Availability

Bulk RNA-seq FASTQ files are deposited in the NCBI Gene Expression Omnibus (GEO, Accession #GSE333280). Reanalysis of snRNA-seq/snATAC-seq multiomic datasets was performed on a deposited dataset in GEO (Accession #GSE294559). R-scripts used for Bulk RNA-seq differential analysis and snRNA-seq/snATAC-seq multiomic analysis can be found on GitHub at https://github.com/kthapa101/siOvol1-bulkseq-analysis and https://github.com/mbaddoo/CRC-endothelial-epithelial-multiome respectively.

---

**Supplementary Table 1. Differentially-expressed genes (DEGS) in si-Ovol1 EC (vs. si-Ctrl EC).** A list of DEGs that exhibited significant (adjusted p-value <0.05) fold expression change vs. control EC.

**Supplementary Table 2. Significant GO Terms in si-Ovol1 EC (vs. si-Ctrl EC) by PANTHER.** A list of significant GO terms identified from the DEG list in **Supplementary Table 1**.

**Supplementary Table 3. Angiogenesis-Associated DEGs in si-Ovol1 EC (vs. si-Ctrl EC).** A list of significant DEGs driving detection of angiogenesis-related GO terms in **Figure 2**.

## Supplementary Figures

**Supplemental Figure 1.**
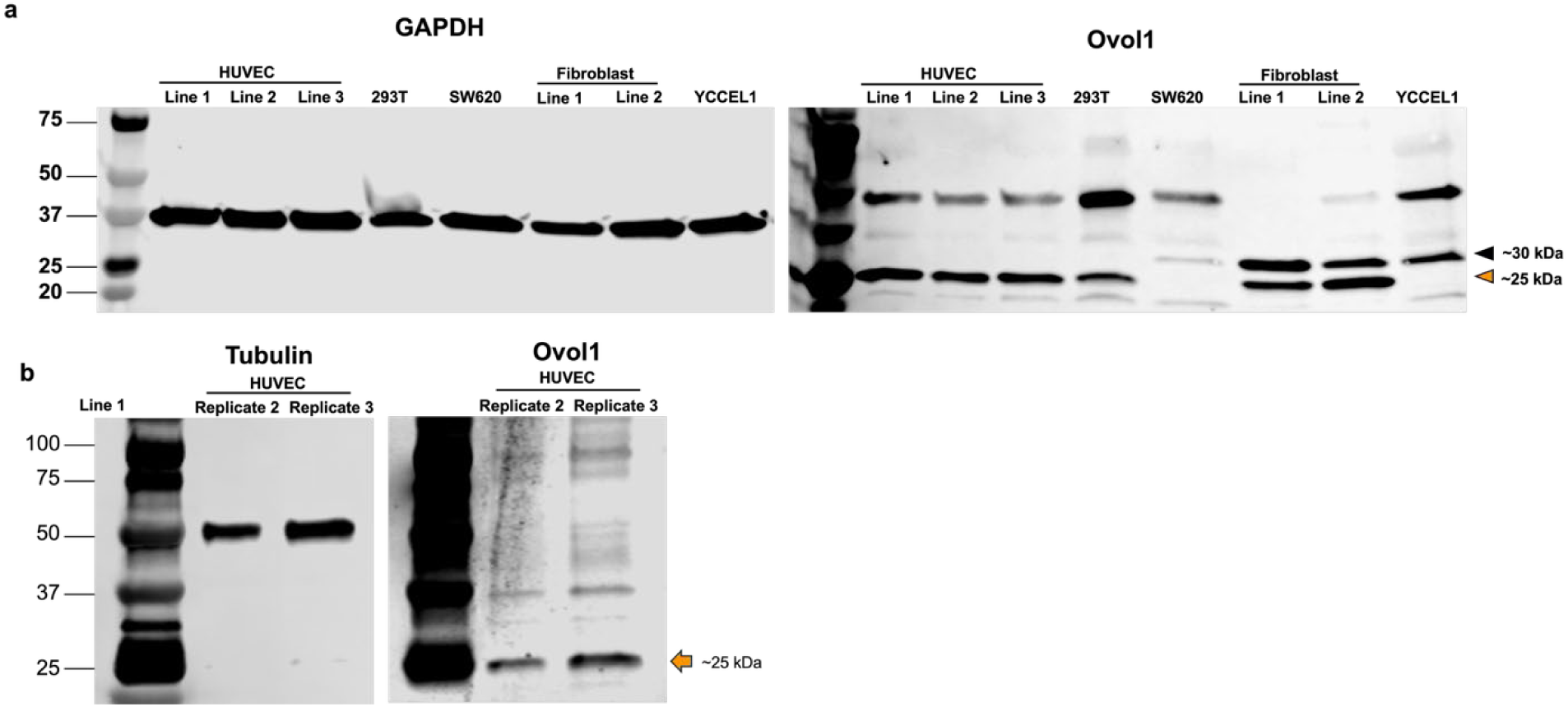
Expanded Western blots for Ovol1 expression. **a)** Protein was isolated from cells cultured in 2D monolayer and probed by Western blot for Ovol1 (or GAPDH as a loading control). Bands were detected at ∼37kDa for Tubulin. Ovol1 was detected at ∼25 kDa in HUVECs and 293T cells, ∼30 kDa in SW620 and YCCEL1 cells, and at both ∼25 and ∼30 kDa in Fibroblasts. **b)** Additional replicates suggesting ∼25kDa Ovol1 band in HUVECs lysate.

**Supplementary Figure 2.**
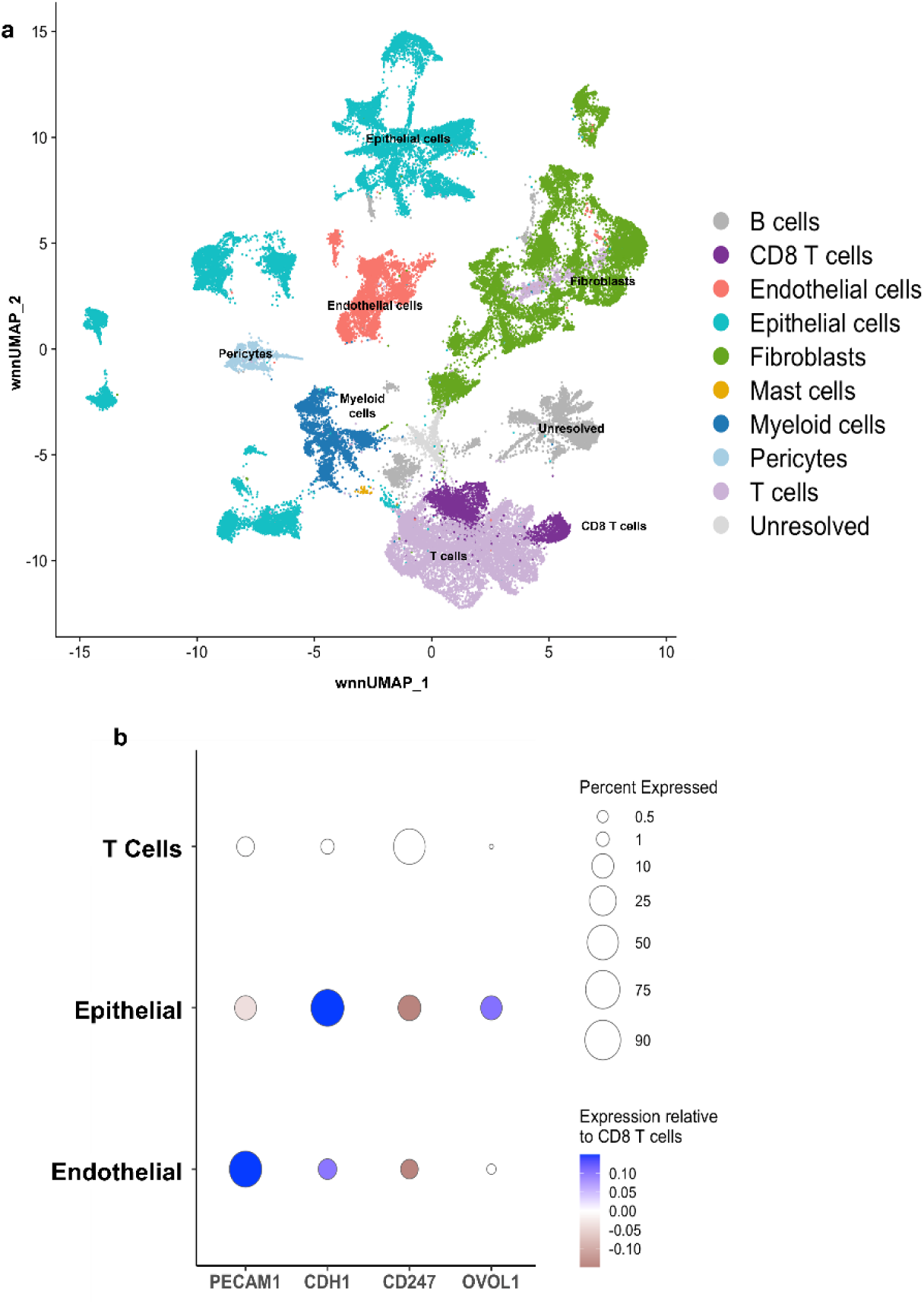
Ovol1 is endogenously expressed at low levels in endothelial cells. **a)** In reanalysis of single-nuclei multiomic analysis of healthy colon tissue [21] (GEO GSE294559), we identify 9 cell populations based on marker expression, including endothelial and epithelial cells. Endothelial cell cluster was identified by expression of endothelial-specific PECAM1 while epithelial cell cluster was identified by expression of epithelial marker CDH1. Both populations were compared against the T-cell population, which was identified by expression of marker CD247 and is negative for Ovol1. In comparison to T-cells, Ovol1 transcript was detectible above baseline in both endothelial and epithelial cell clusters, with higher expression evident for epithelial cells compared to endothelial cells.

**Supplementary Figure 3.**
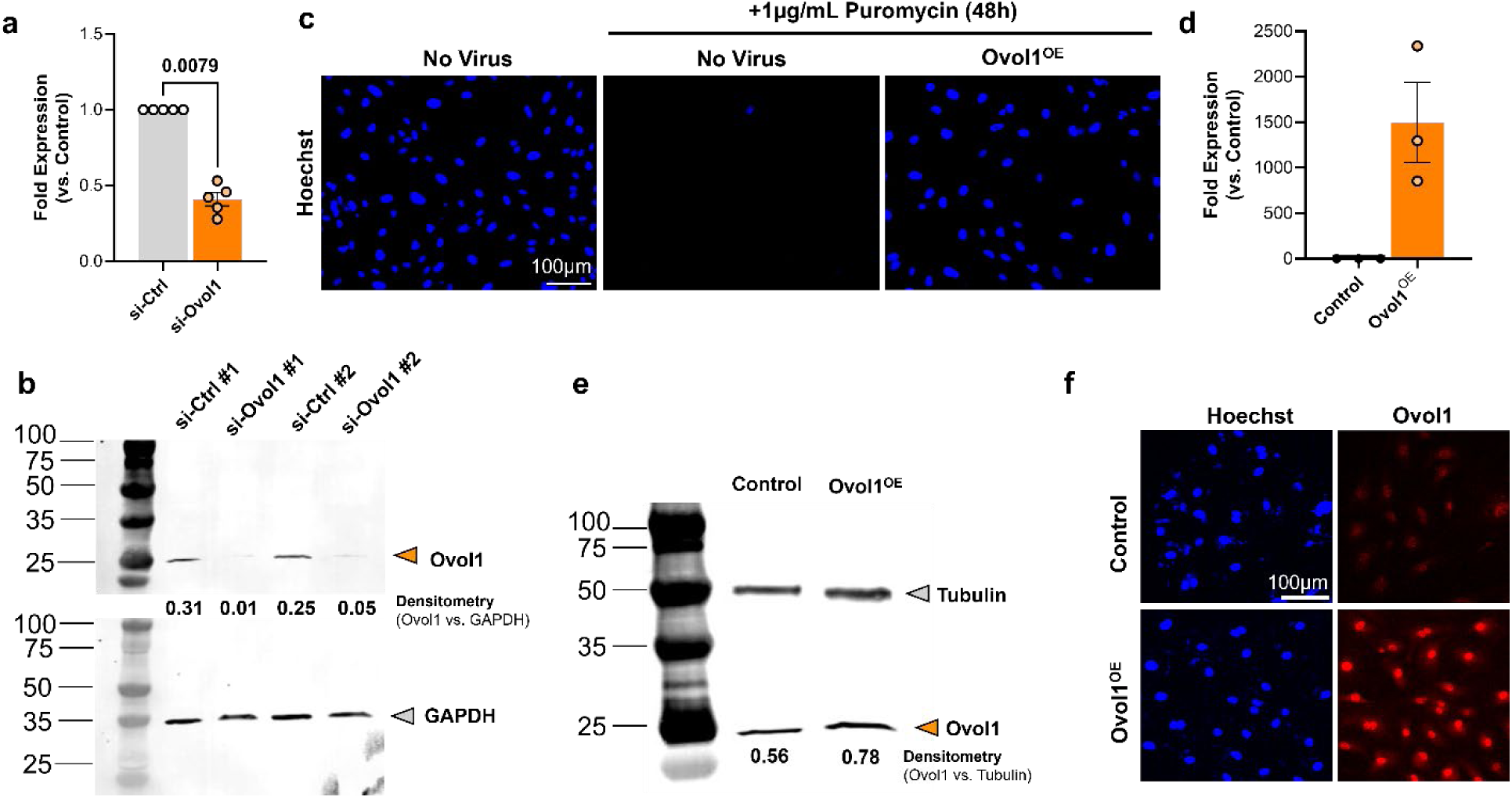
Validation of Ovol1 knockdown and overexpression tools. **a)** Silencing RNA knockdown eliminates ∼55% of endogenous Ovol1 mRNA and **b)** reduces Ovol1 protein expression. (Statistics: Wilcoxon rank-sum test). **c)** Exposure to Ovol1 overexpression (Ovol1^OE^) lentivirus (MOI ≈ 1) confers puromycin resistance on HUVEC. Ovol1 **d)** mRNA and **e)** protein levels are increased in Ovol1^OE^ EC compared to EC transduced with control (eGFP) lentivirus. **f)** Constitutively expressed Ovol1 is localized to the nucleus in Ovol1^OE^ EC.

**Supplementary Figure 4.**
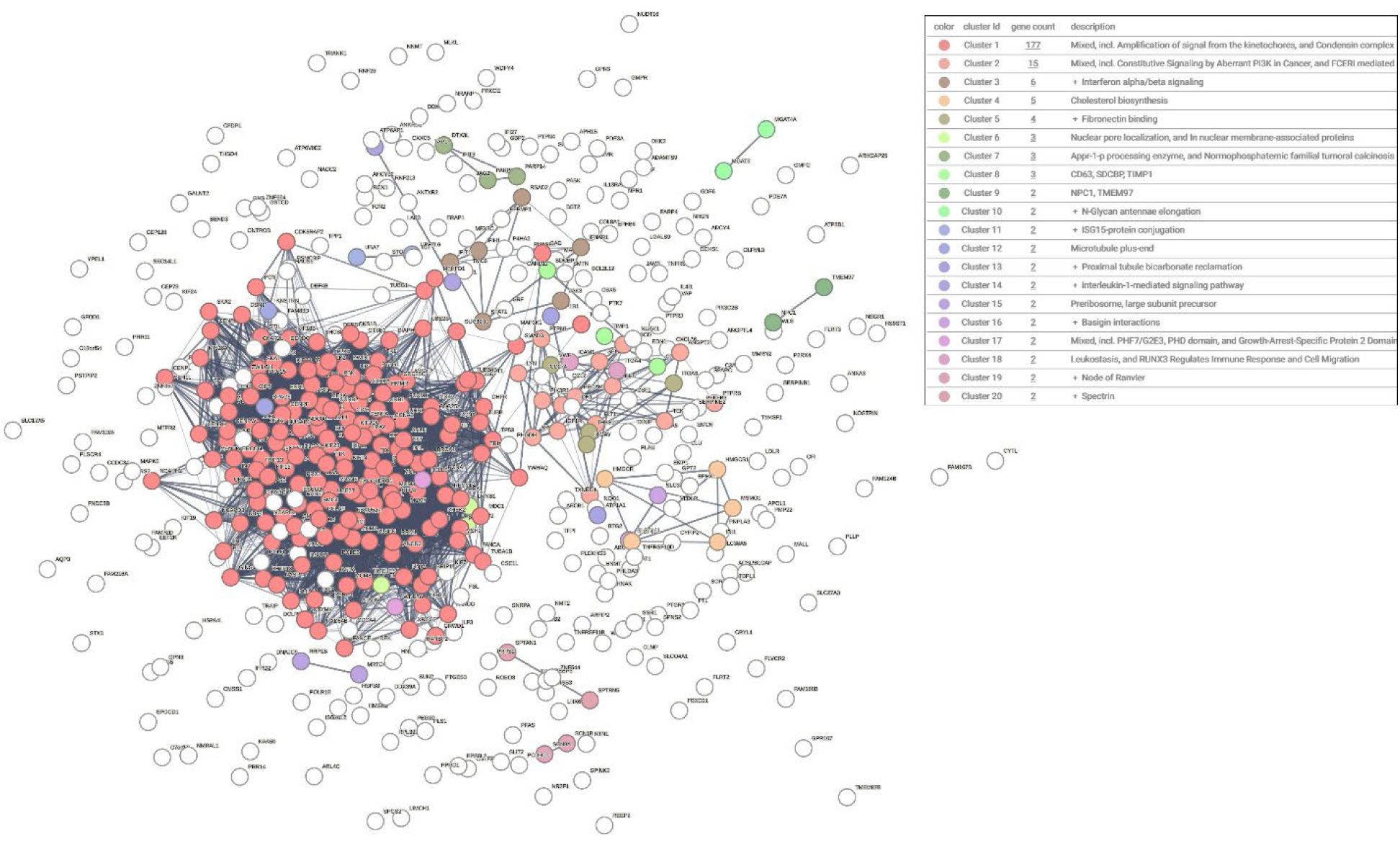
Shared Ovol1 and Slug genes identify interaction clusters related to cell proliferation. Interaction network analysis of the 579 genes identified as regulated by both Ovol1 and Slug in **Figure 2d** identifies 20 interaction clusters. Of these, the largest two clusters are associated with cell cycle and cell proliferation (Cluster 1, red), and associated pro-growth pathways such as PI3K (Cluster 2, orange).

**Supplementary Figure 5.**
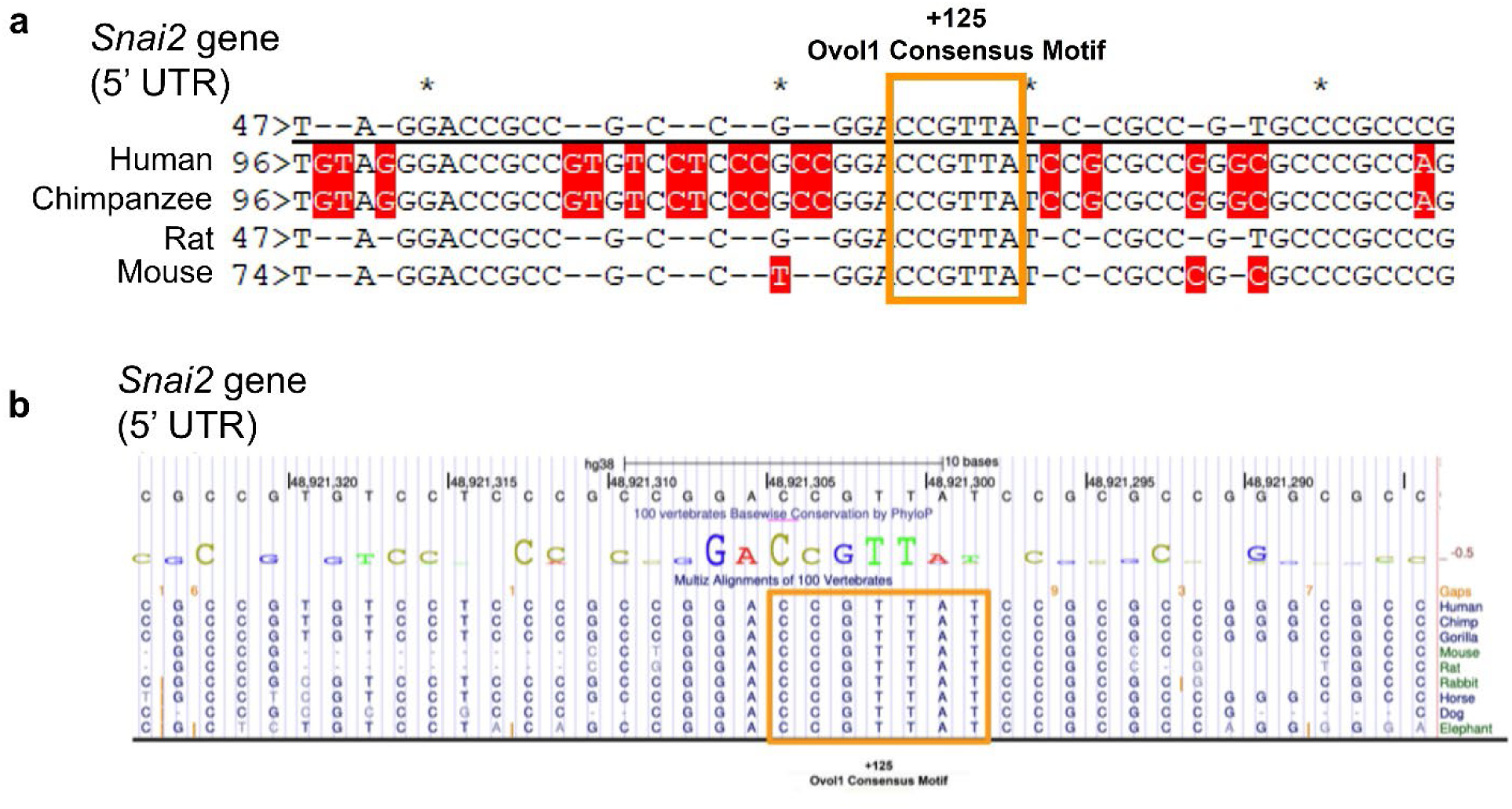
Conservation of predicted Ovol1 consensus site in Slug gene. **a)** Alignment of the 5’ untranslated region (UTR) of the Snai2 gene from human (Genbank Acc. No: NG_012130.1), chimpanzee (Genbank Acc. No: XM_016959457.3), rat (Genbank Acc. No: NM_013035.2) and mouse (Genbank Acc. No: NM_011415.3) reveals that the predicted Ovol1 consensus sequence motif CCGTTA (nucleotide positions 125-130 in humans) is fully conserved across all four species. **b)** Using the USCS Genome Browser,[27] we separately also identified strong conservation of the +125 Ovol1 binding site in the Slug gene across 100 compared vertebrate species.

**Supplementary Figure 6.**
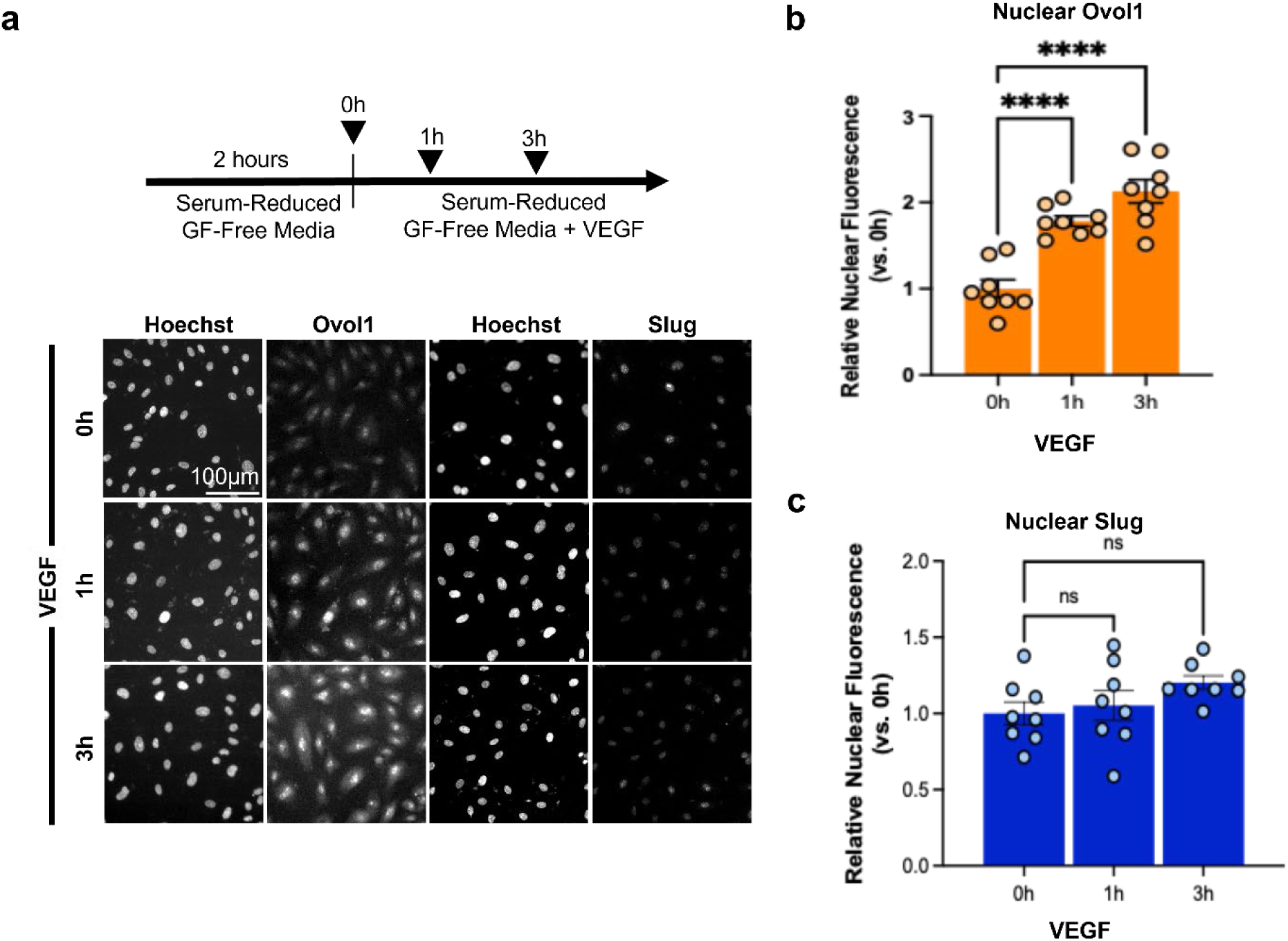
Nuclear Ovol1 is upregulated by VEGF prior to upregulation of nuclear Slug. **a)** HUVEC monocultures were serum- and growth-factor starved for 2h, followed by treatment with 10ng/mL VEGF for up to 6h following by cell fixation and immunofluorescence for Ovol1 or Slug. Cell nuclei were counter-stained with Hoechst 333452. **b)** Mean nuclear fluorescence for Ovol1 was low at 0h and increased within 1h (vs. 0h) of VEGF stimulation. **c**) By contrast, nuclear Slug was not significantly increased within this timeframe (8-9 technical replicates per group, Statistics: 1-way ANOVA with post hoc t-test adjusted for multiple comparisons).

**Supplementary Figure 7.**
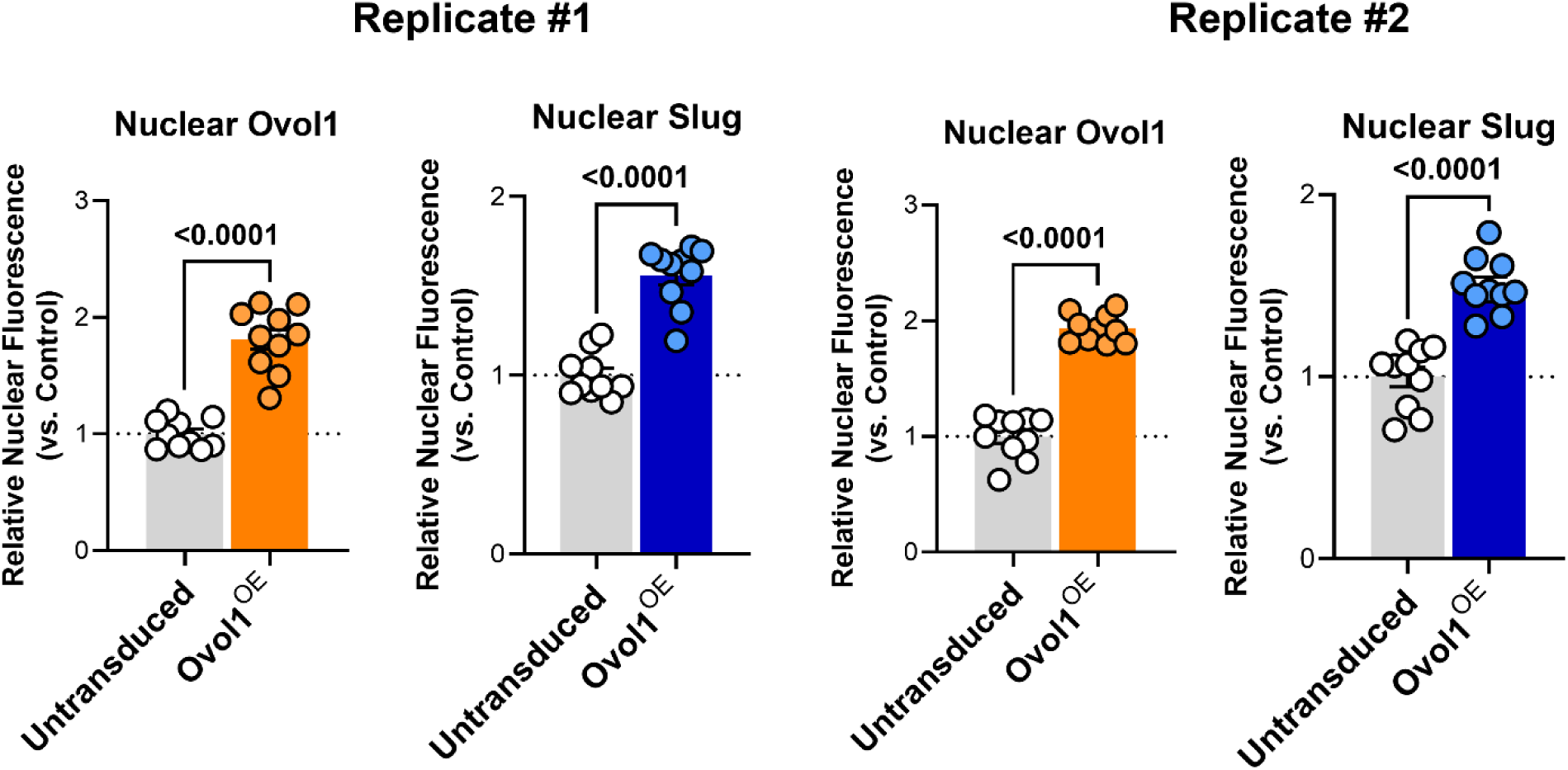
Slug is increased in Ovol1-overexpressing endothelial cells compared to untransduced controls. In two independent replicates (each technically sampled 10 times), nuclear Ovol1 was increased in Ovol1^OE^ endothelial cells compared to untransduced endothelial cells, alongside a corresponding significant upregulation of nuclear Slug.

**Supplementary Figure 8.**
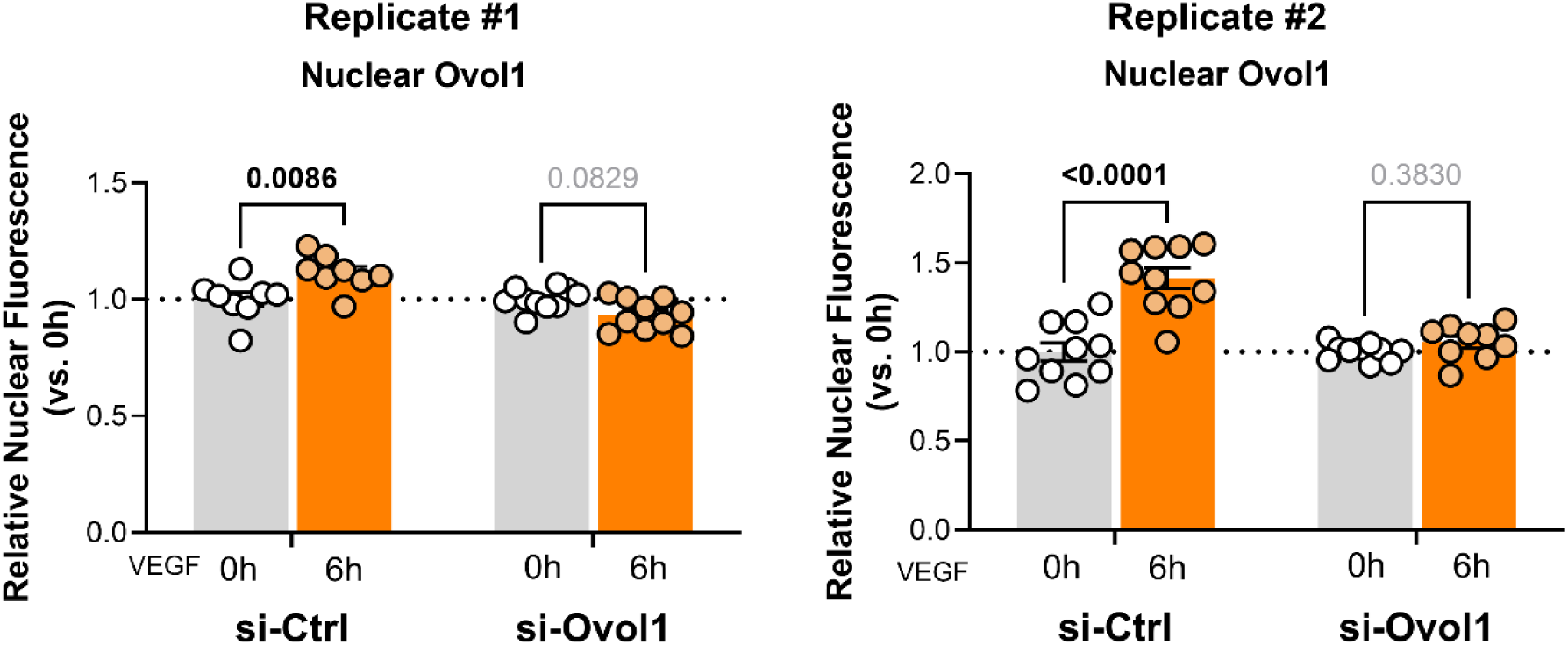
si-Ovol1 prevents VEGF-induced upregulation of Ovol1. In two independent replicates (each technically sampled 8 times), VEGF stimulation of endothelial cells increases nuclear Ovol1 at 6h, but this upregulation is prevented by pre-treatment with si-Ovol1.

**Supplemental Figure 9.**
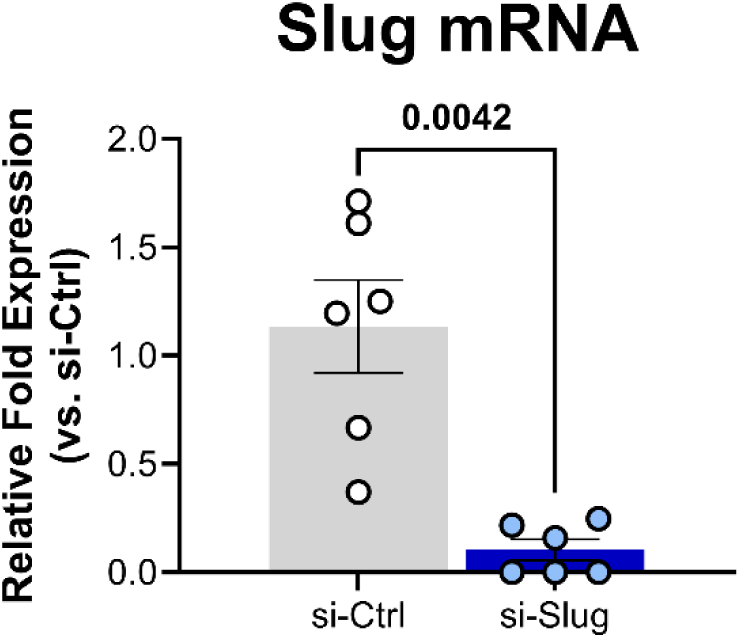
Validation of silencing RNA knockdown of endogenous Slug. HUVEC monocultures transfected with si-Slug expressed significantly less endogenous Slug mRNA compared to cells treated with non-targeting control siRNA (i.e., si-Ctrl). (n=6 si-Ctrl, n=6 si-Slug independent replicates, Statistics: Unpaired Students’ t-test).

**Supplementary Figure 10.**
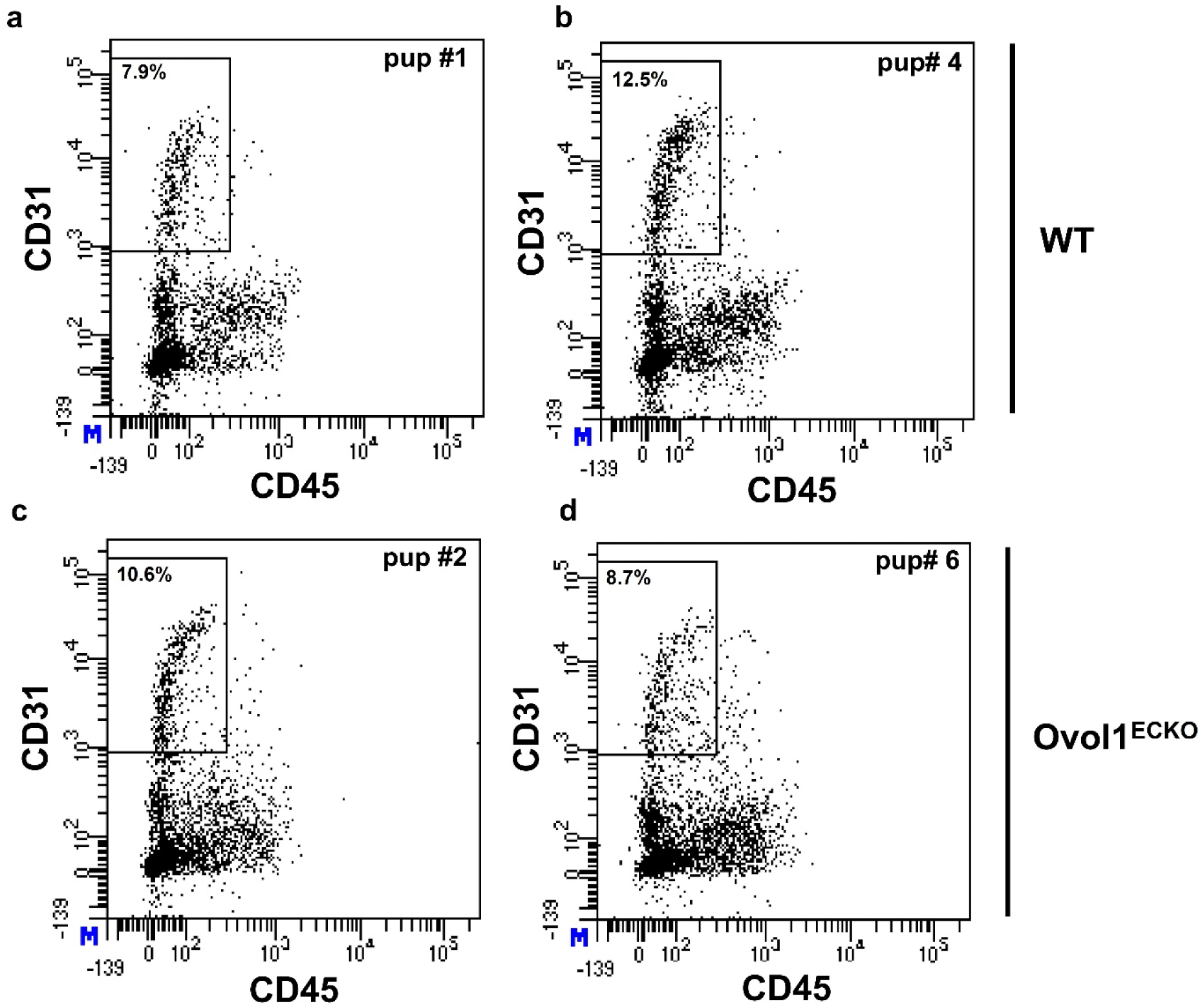
FACS sorting of murine lung endothelial (CD31⁺/CD45⁻) cells for wild-type and Ovol1^ECKO^ mice. Single-cell suspensions were generated from lung tissue harvested from P15 Cre-negative wild-type (WT) and Cre-positive Ovol1^ECKO^ mouse littermates. FACS was used to sort for endothelial (CD31^+^/CD45^−^) cells. FACS scatter plots show gating for 50,000 cells for **a-b)** WT and **c-d)** Ovol1^ECKO^ mouse lines and reveals similar endothelial cell proportions for each genotype.

**Supplementary Figure 11.**
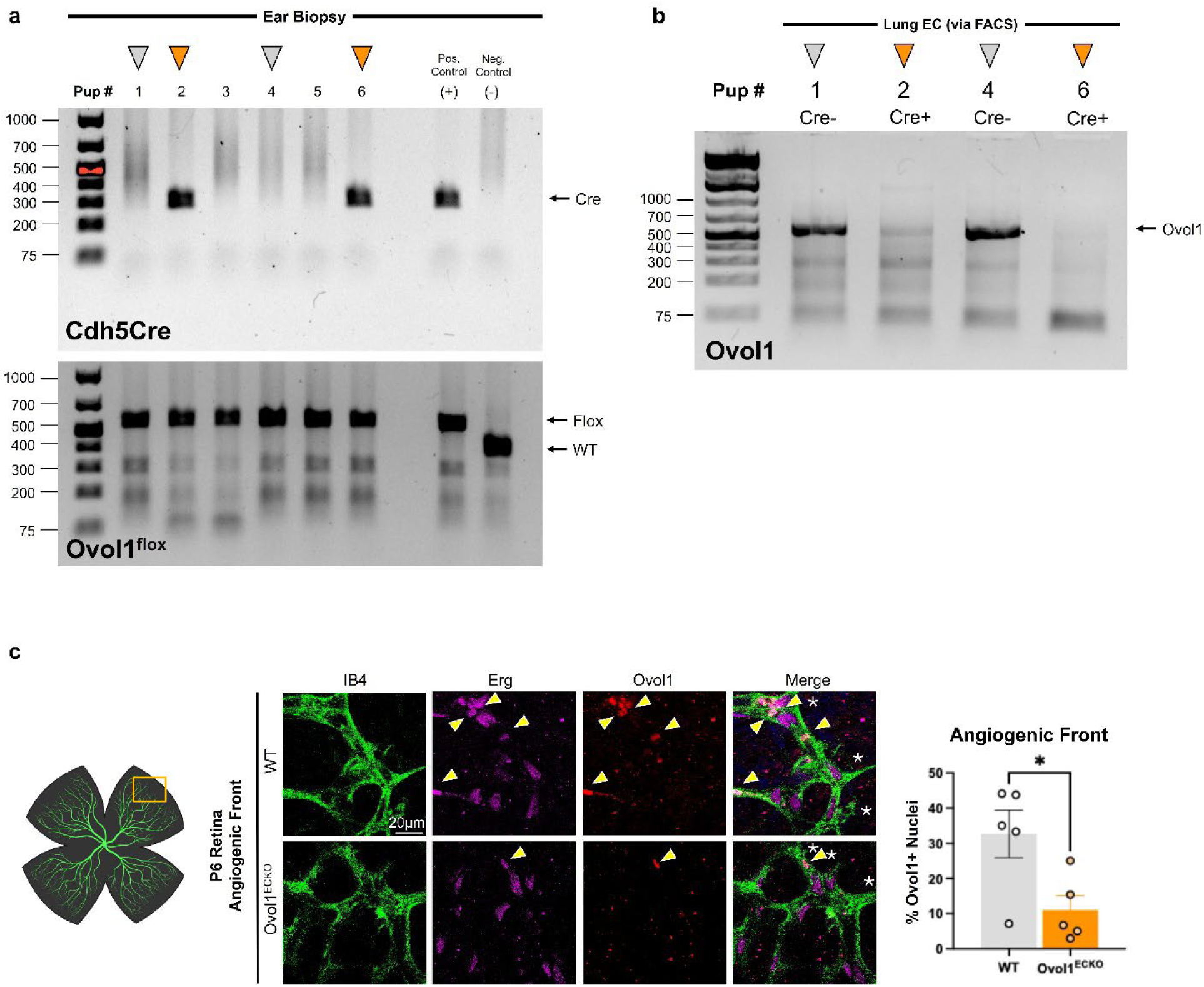
Validation of endothelial Ovol1 knockout in Ovol1^ECKO^ mouse line. **a)** 6 post-natal day 15 littermates from the Ovol1^ECKO^ line were biopsied and genotyped by PCR for Cre and Ovol1^flox^ status, which identified that all mice were homozygous for Ovol1^flox^ and that mice #2 and #6 were Cre-positive (orange arrowhead), consistent with the breeding scheme that generated this litter. **b)** End-point PCR was performed on genomic DNA from ∼500,000 FACS-sorted of lung (CD31^+^/CD45^−^) EC. The Ovol1 gene was robustly amplified in Cre-negative mice (#1 and #4, grey arrowhead) but not in Cre-positive mice (#2 and #6, orange arrowhead) indicating ablation of the Ovol1 gene in vascular endothelium. **c)** The neonatal retinal vasculature of wild-type (i.e., Cre-;Ovol1^flox/flox^) and Ovol1^ECKO^ (i.e., Cre+; Ovol1^flox/flox^) mice harvested at post-natal day 6 and stained by immunofluorescence for Ovol1. Vessels were counterstained with the endothelial-specific marker IB4 and the endothelial nuclei-specific marker Erg. The number of Ovol1+ EC nuclei were counted in the angiogenic front. Frequency of Ovol1+ EC nuclei was significantly reduced by ∼60-70% in Ovol1^ECKO^ mice, indicating efficient (but incomplete) penetrance of knockout in these mice. (Statistics: Students’ t-test).

**Supplementary Figure 12.**
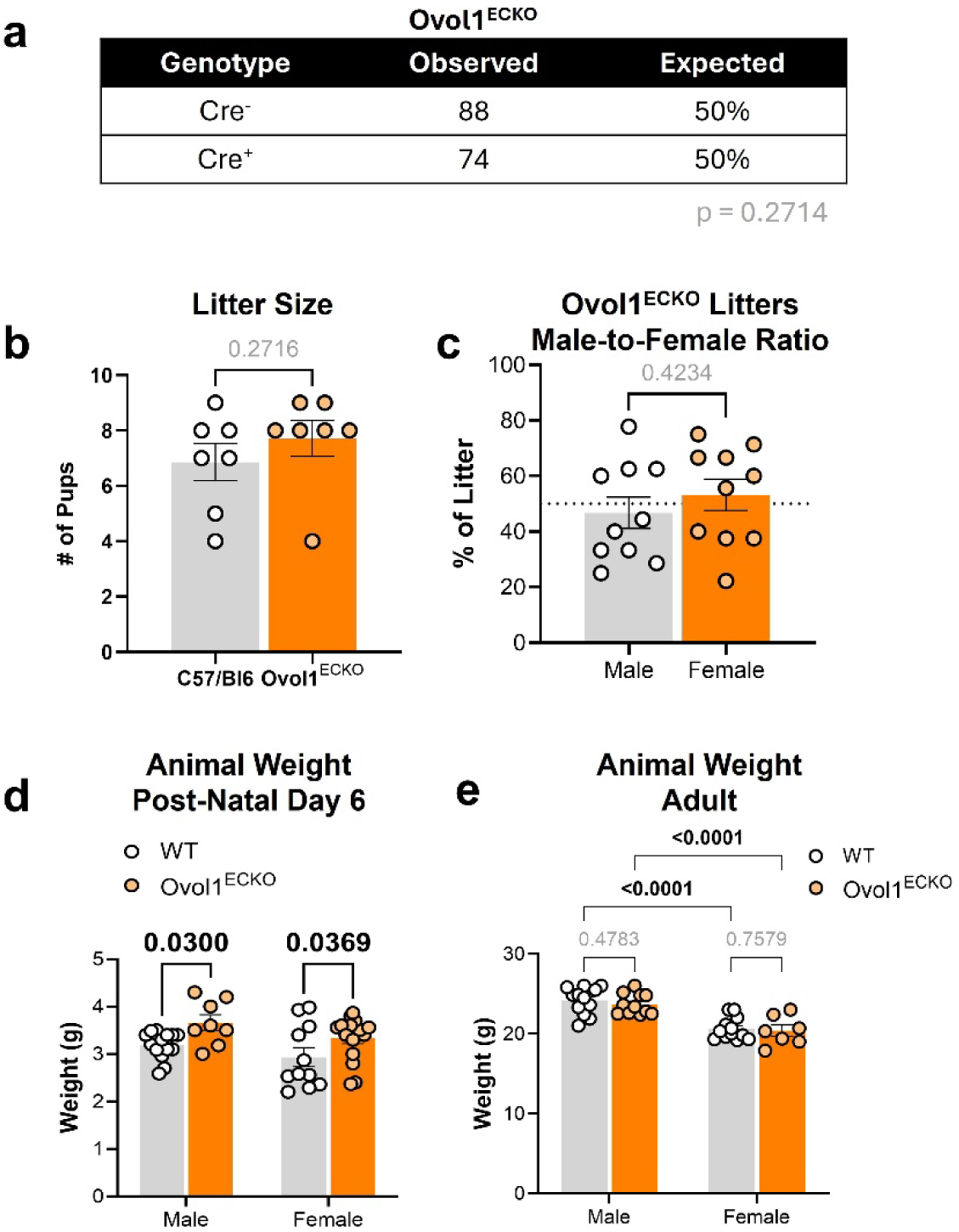
Vital characteristics of Ovol1^ECKO^ mice. **a)** Cre+ and Cre^−^ mice were born at expected Mendelian ratios in Ovol1^ECKO^ mouse line (Statistics: Chi square test, litters from 9 independent breeding pairs). **b)** Average litter size of Ovol1^ECKO^ breeding pairs was similar to that of C57/Bl6 wild-type mice (Statistics: Wilcoxon rank-sum test). **c)** Male-to-female ratio was approximately equal across multiple Ovol1^ECKO^ litters (Wilcoxon rank-sum test). **d)** Ovol1^ECKO^ pups are slightly larger than their wild-type littermates at post-natal day 6 (Statistics: 2-way ANOVA with post-hoc adjustment for multiple comparisons), but **e)** no differences in animal weight were detected between wild-type vs. Ovol1^ECKO^ mice by 20 weeks of age. As expected, male mice were slightly heavier than female mice regardless of genotype (Statistics: 2-way ANOVA with post-hoc t-test).

**Supplementary Figure 13.**
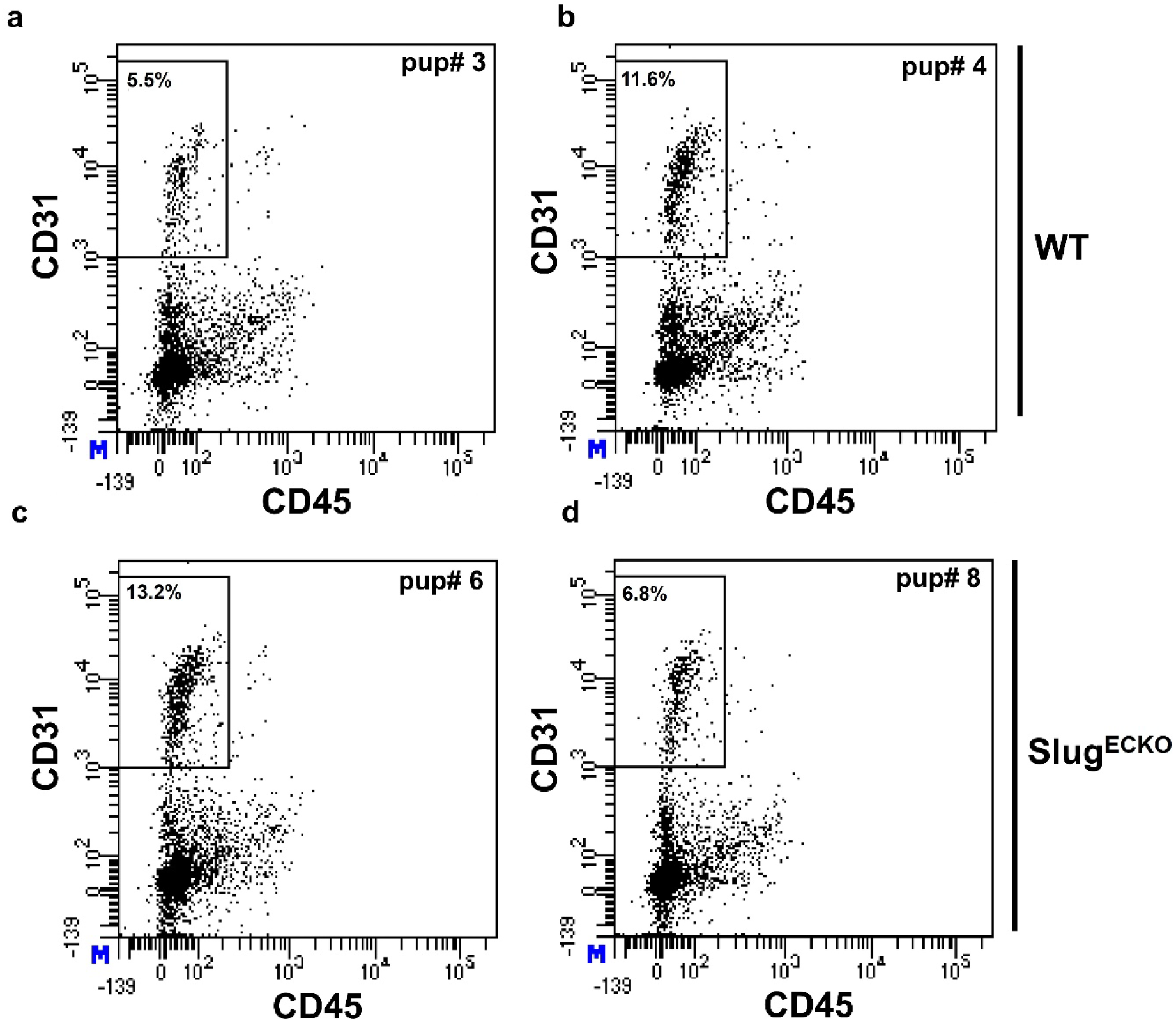
FACS sorting of murine lung endothelial (CD31⁺/CD45⁻) cells for wild-type and Slug^ECKO^ mice. Single-cell suspensions were generated from lung tissue harvested from P15 Cre-negative wild-type (WT) and Cre-positive Slug^ECKO^ mouse littermates. FACS was used to sort for endothelial (CD31^+^/CD45^−^) cells. FACS scatter plots show gating for 50,000 cells for **a-b)** WT and **c-d)** Slug^ECKO^ mouse lines and reveals similar endothelial cell proportions for each genotype.

**Supplementary Figure 14.**
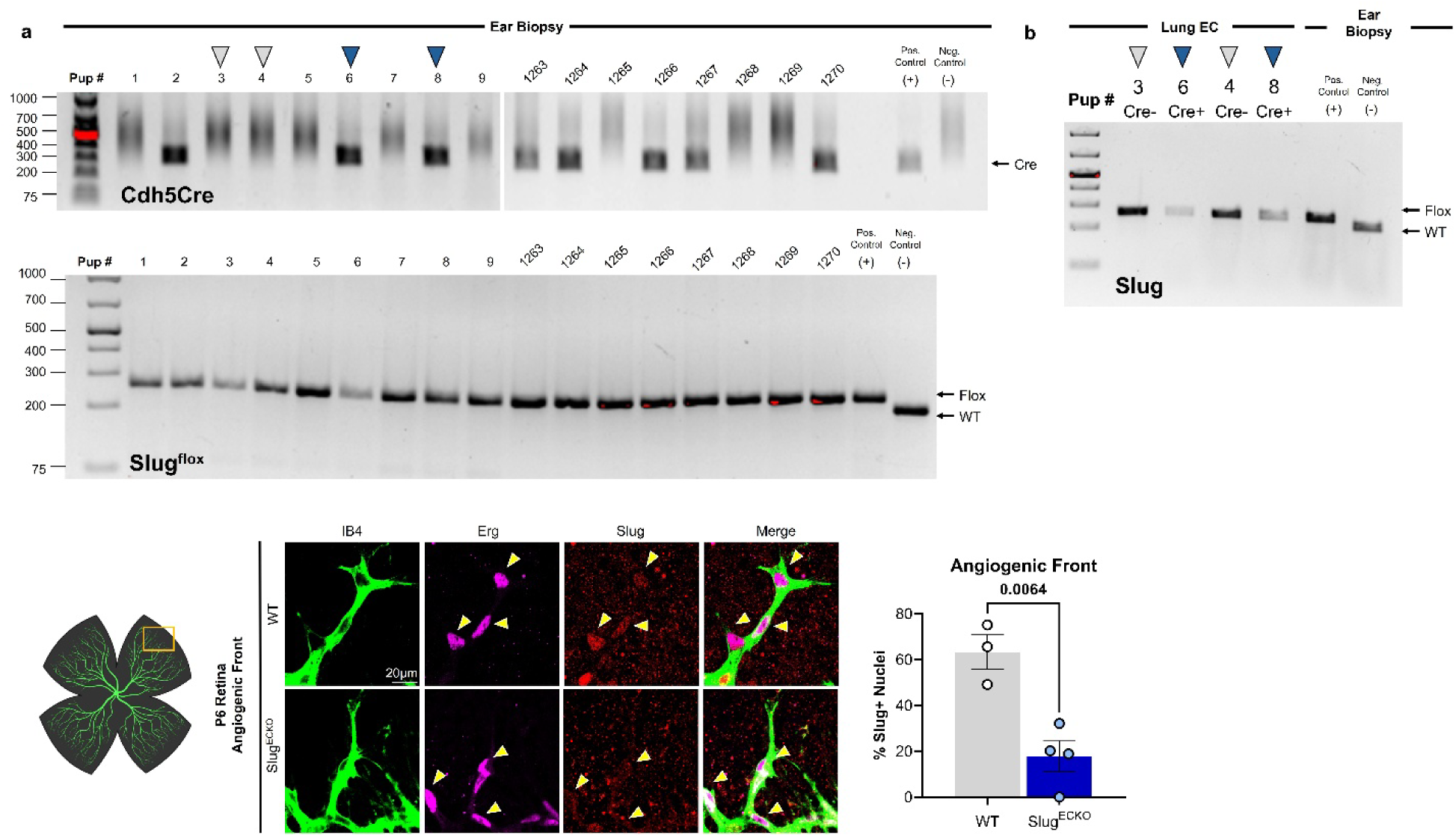
Validation of endothelial Slug knockout in Slug^ECKO^ mouse line. **a)** 9 post-natal day 15 littermates from the Slug^ECKO^ line were biopsied and genotyped by PCR for Cre and Slug^flox^ status, which identified that all mice were homozygous for Slug^flox^ and that mice #2, 6, and #8 were Cre-positive (blue arrowhead), consistent with the breeding scheme that generated this litter. In parallel, 8 adult Slug^ECKO^ were genotype, and positive and negative control samples were run to facilitate identification of bands. **b)** End-point PCR was performed on genomic DNA from ∼500,000 FACS-sorted of lung (CD31^+^/CD45^−^) EC. The Slug gene was robustly amplified in Cre-negative mice (#3 and #4, grey arrowhead) but was poorly amplified in Cre-positive mice (#6 and #8, blue arrowhead) indicating endothelial-specific knockout of Slug with efficient (but incomplete) penetrance. **c)** The neonatal retinal vasculature of wild-type (i.e., Cre-;Slug^flox/flox^, n= 3) and Slug^ECKO^ (i.e., Cre+; Slug^flox/flox^, n= 4) mice harvested at post-natal day 6 and stained by immunofluorescence for Slug. Vessels were counterstained with the endothelial-specific marker IB4 and the endothelial nuclei-specific marker Erg. The number of Slug+ EC nuclei were counted in the angiogenic front. Frequency of Slug+ EC nuclei was significantly reduced by ∼70% in Slug^ECKO^ mice. (Statistics: Unpaired Students’ t-test).

**Supplementary Figure 15.**
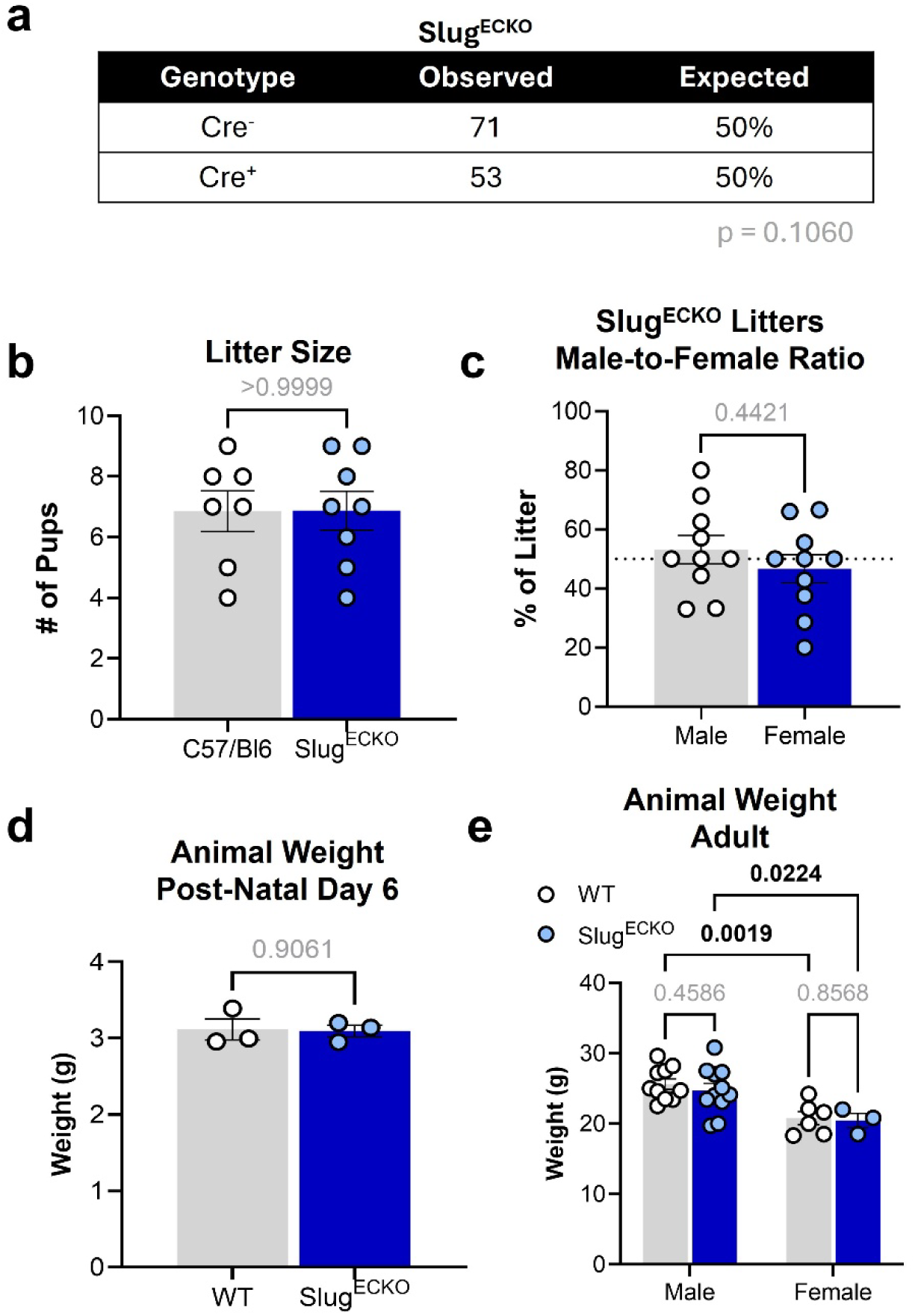
Vital characteristics of Slug^ECKO^ mice. **a)** Cre+ and Cre^−^ mice were born at expected Mendelian ratios in Slug^ECKO^ mouse line (Statistics: Chi square test, litters from 8 independent breeding pairs). **b)** Average litter size of Slug^ECKO^ breeding pairs was similar to that of C57/Bl6 wild-type mice (same control data as graphed in **Supplementary Figure 13b**, Statistics: Wilcoxon rank-sum test). **B)** Male-to-female ratio was approximately equal across multiple Slug^ECKO^ litters (Statistics: Wilcoxon rank-sum test). **c)** No difference in animal weight was observed between Slug^ECKO^ mice and wild-type littermates at post-natal day 6 (males and females combined, Statistics: Welch’s t-test). **d)** No genotype-related differences in overall animal weight were observed at 20 weeks of age, although male mice were heavier than female mice regardless of genotype as is typically expected (Statistics: 2-way ANOVA with post-hoc adjustment for multiple comparisons).

**Supplementary Figure 16.**
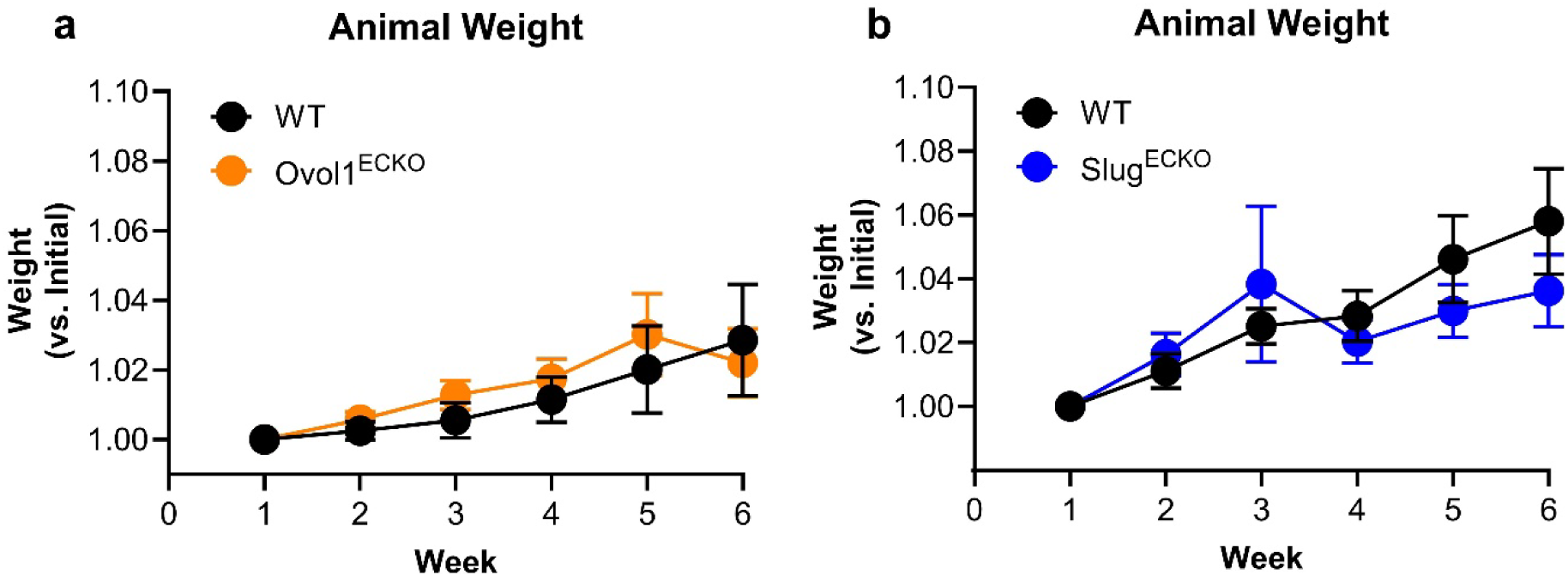
Relative animal weight during tumor growth in colorectal cancer syngraft model. No differences were observed across genotype in the relative increase in animal weight for both **a)** Ovol1^ECKO^ (n=13 WT, n=21 Ovol1^ECKO^) or **b)** Slug^ECKO^(n=14 WT, n=25 Slug^ECKO^) mice following subcutaneous injection of CMT93 colorectal cancer syngrafts. In both graphs, there is a statistically significant increase in animal weight over 6 weeks following tumor engraftment.

**Supplementary Figure 17.**
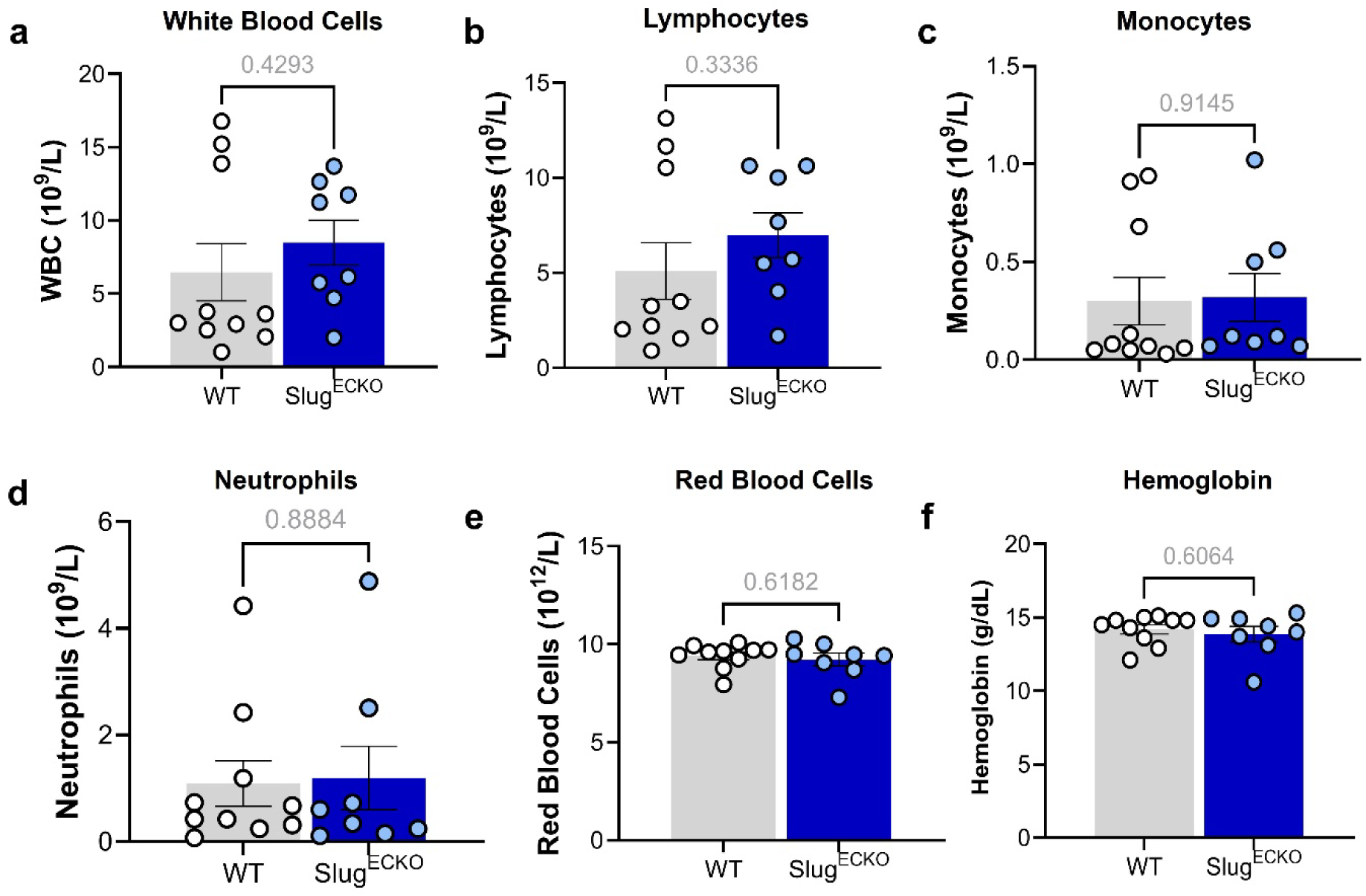
Hematological profiles of WT vs Slug^ECKO^ mice. No statistically significant differences were observed between Cre-wild-type and Cre+ Slug^ECKO^ littermates for blood cell counts, including for **a)** white blood cells, **b)** lymphocytes, **c)** monocytes, **d)** neutrophils, **e)** or red blood cells. **f)** Hemoglobin levels were also comparable across mouse genotypes. (Statistics: Unpaired Students’ t-test).

**Supplementary Figure 18.**
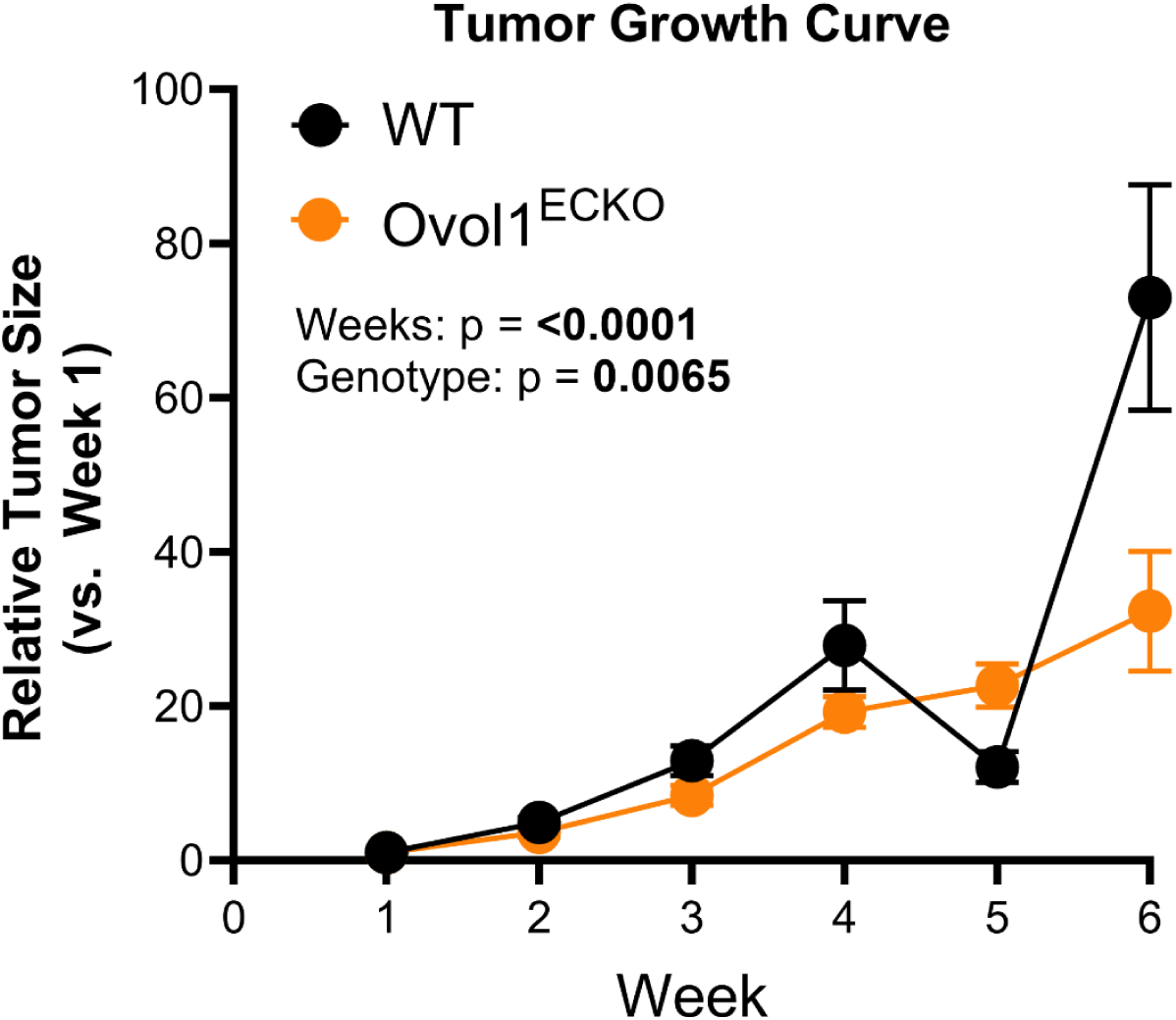
Growth of MC38 colorectal cancer syngrafts are also altered in Ovol1^ECKO^. MC38 murine adenocarcinoma cells were injected into Ovol1^ECKO^, and tumor growth rate was observed to be significantly different compared to tumors engrafted in wild-type (Cre^−^) littermates (n=20 Wild-type, n=11 Ovol1^ECKO^ mouse littermates, Statistics: 2-way ANOVA).

**Supplementary Figure 19.**
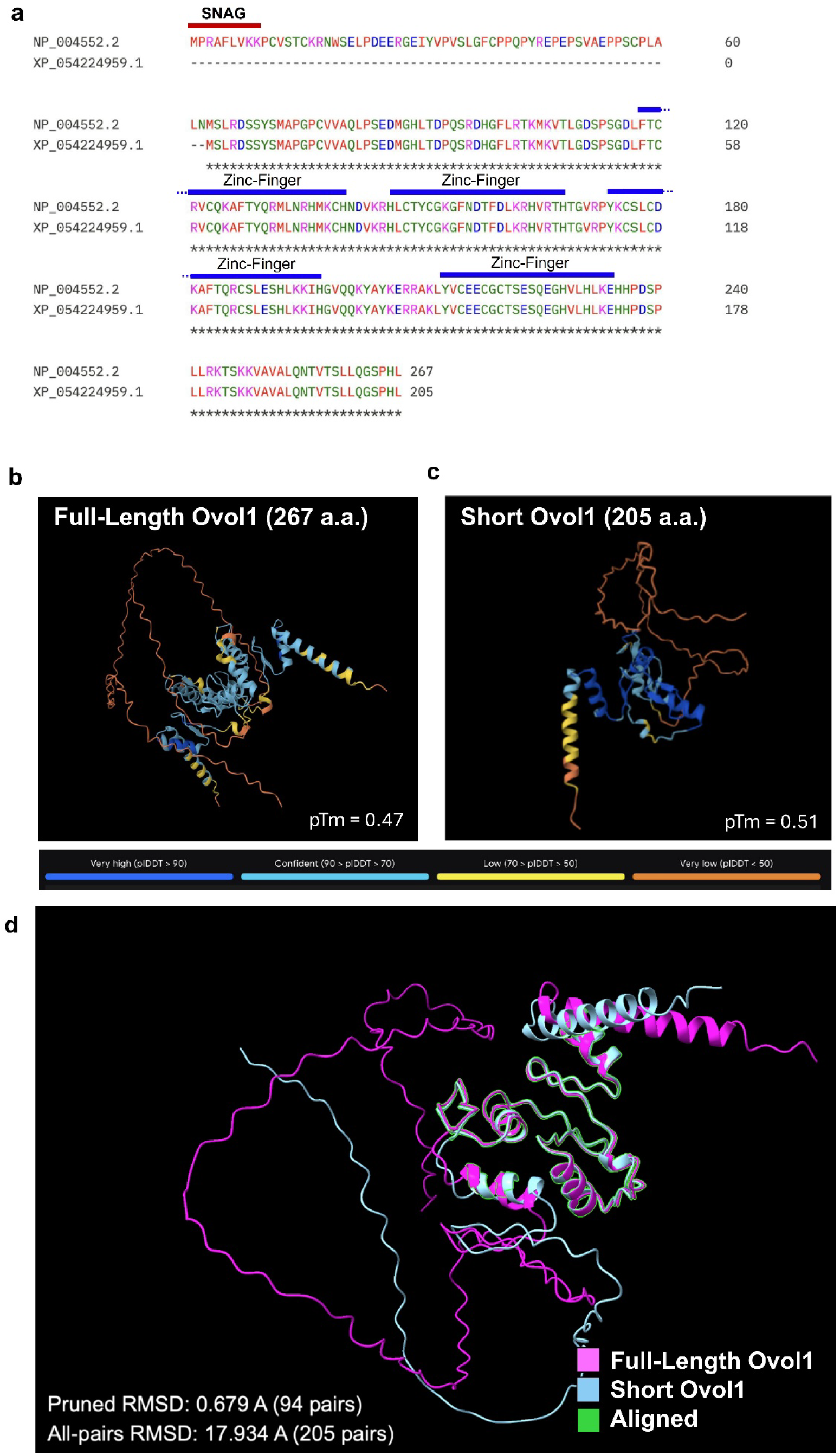
Structural comparison of full-length and predicted N-terminal truncated Ovol1 variants. **a)** Sequence alignment of full-length Ovol1 (Genbank NP_004552.2) vs. a predicted N-terminal truncated variant from RefSeq (Genbank XP_054224959.1). **b-c)** Structural prediction of Ovol1 variants (full-length Genbank NP_004552.2, short Genbank XP_054224959.1). **d)** Subsequent comparison of structures in **b-c** using ChimeraX reveals that the truncated variant lacks N-terminal SNAG domain and unstructured region, but aligns strongly with full-length Ovol1 for conserved zinc-finger binding domains closer to the C-terminus suggesting preservation of DNA-binding capacity.

